# Parallel genetic evolution and speciation from standing variation

**DOI:** 10.1101/368324

**Authors:** Ken A. Thompson, Matthew M. Osmond, Dolph Schluter

**Affiliations:** Biodiversity Research Centre and Department of Zoology, University of British Columbia, Vancouver, Canada; Center for Population Biology, University of California, Davis, USA

**Keywords:** Fisher’s geometric model, ecological speciation, mutation-order speciation, simulation, mathematical theory, convergent evolution, segregation variance, hybrid breakdown, parallel evolution

## Abstract

Adaptation often proceeds via the sorting of standing variation, and natural selection acting on pairs of populations is a quantitative continuum ranging from parallel to divergent. Yet, it is unclear how the extent of parallel genetic evolution during adaptation from standing variation is affected by the difference in the direction of selection between populations. Nor is it clear whether the availability of standing variation for adaptation affects progress toward speciation in a manner that depends on the difference in the direction of selection. We conducted a theoretical study investigating these questions and have two primary findings. First, the extent of parallel genetic evolution between two populations is expected to rapidly decline as the difference in their directions of selection increases from fully parallel toward divergent, and this decline occurs more rapidly in organisms with greater trait ‘dimensionality’. This rapid decline results because seemingly small differences in the direction of selection cause steep reductions in the fraction of alleles that are beneficial in both populations. For example, populations adapting to optima separated by an angle of 33° have only 50% of potentially beneficial alleles in common (for a case of five trait ‘dimensions’). Second, we find that adaptation from standing variation leads to higher ecologically-dependent hybrid fitness under parallel selection, relative to when adaptation is from new mutation only. This occurs because genetic parallelism based on standing variation reduces the phenotypic segregation variance in hybrids when parents adapt to similar environments. In contrast, under divergent selection, the pleiotropic effects of alternative alleles fixed from standing variation change the major axes of phenotypic variation in hybrids and reduce their fitness in parental habitats. We conclude that adaptation from standing genetic variation is expected to slow progress toward speciation via parallel natural selection and can facilitate progress toward speciation via divergent natural selection.

**Impact summary:** It is increasingly clear that much of adaptation, especially that which occurs rapidly, proceeds from the sorting of ancestral standing variation rather than complete reliance on *de novo* mutation. In addition, evolutionary biologists are increasingly embracing the fact that the difference in the direction of natural selection on pairs of populations is a quantitative continuum ranging from completely parallel to completely divergent. In this article, we ask two questions. First, how does the degree of genetic parallelism—here, adaptation using the same alleles in allopatric populations—depend on the differences in the direction of natural selection acting on two populations, from parallel (0°) to divergent (180°)? And second, how does adaptation from standing variation affect progress toward speciation, and does its effect depend on the direction of natural selection? We develop theory to address these questions. We first find that very small differences in the direction of selection (angle) can largely preclude genetic parallelism. Second, we find that adaptation from standing variation has implications for speciation that change along the continuum from parallel to divergent selection. Under parallel selection, high genetic parallelism causes inter-population hybrids to have high mean fitness when their parents adapt from standing variation. As selection tends toward divergent, adaptation from standing variation is less beneficial for hybrid fitness and under completely divergent selection causes inter-population hybrids to have lower mean fitness than when adaptation was from new mutation alone. In sum, our results provide general insight into patterns of genetic parallelism and speciation along the continuum of parallel to divergent natural selection when adaptation is from standing variation.

## Introduction

In recent years, two general features of evolution by natural selection have become increasingly established. First, adaptation often proceeds largely via the reassortment of ancestral standing variation rather than via complete reliance on *de novo* mutations (Barrett and Schluter 2008). And second, variation in the direction of natural selection acting on pairs of populations is best represented by a quantitative continuum ranging from parallel selection—favouring identical phenotypes—to divergent selection—favouring distinct phenotypes — rather than falling into discrete ‘parallel’ or ‘divergent’ bins (Bolnick et al. 2018). It is unclear, however, how the extent of parallel genetic evolution—use of the same alleles during adaptation—might depend on the difference in the direction of selection experienced by a pair of populations. Whether or not populations undergo parallel genetic evolution has consequences for the evolution of reproductive isolating barriers between them (Schluter and Conte 2009). Yet, it is also unclear whether adaptation from standing variation has implications for speciation that are distinct from when adaptation is from new mutation alone, and whether its effect changes along the continuum from parallel to divergent natural selection.

Adaptation facilitates progress towards speciation when populations evolve reproductive isolating barriers as a by-product. One reason these reproductive isolating barriers might arise is because genetic differences between populations have maladaptive consequences when combined in hybrids (i.e., ‘postzygotic’ isolation), thereby reducing gene flow upon secondary contact. When a pair of populations adapts in response to divergent natural selection, hybrids might have an intermediate or mismatched phenotype and be unfit in both parental environments (Hatfield and Schluter 1999; Arnegard et al. 2014; Cooper et al. 2018). When a pair of populations are subject to parallel selection, they may diverge genetically by chance (Mani and Clarke 1990; Schluter 2009) and hybrids might have novel transgressive phenotypes that are poorly suited to the common parental habitat (Barton 1989). How adaptation from standing variation affects progress toward speciation-by-selection (Langerhans and Riesch 2013) is largely unexplored theoretically.

Adaptation from standing variation is common (Barrett and Schluter 2008) and underlies some of the most spectacular adaptive radiations found in nature (Brawand et al. 2015). Genomic studies in some systems also implicate standing variation as the major source of genetic parallelism in replicate populations colonizing similar environments (Jones et al. 2012; Roesti et al. 2014; Lee and Coop 2017). Previous research has shown that the correlation between selection coefficients of a given allele in each of two populations inhabiting different environments increases with the similarity in the direction of selection (equation 6 in Martin and Lenormand 2015). Given this, we expected the extent of parallel genetic evolution for two populations to decline from a maximum to a minimum value as the angle between the directions of selection between them (*θ*) increases from completely parallel (*θ* = 0°) to completely divergent (*θ* = 180°). Our specific goal was to characterize the pattern of decline in parallelism. We also hypothesized that adaptation from standing variation would reduce the evolution of reproductive isolation under parallel selection because parental populations would fix more of the same alleles and therefore evolve fewer incompatibilities (Schluter 2009). Under divergent selection we hypothesized that populations would fix alternative alleles regardless of whether they were selected from standing variation or new mutation. Therefore, we expected standing variation to have little effect on speciation by divergent selection compared to adaptation from new mutation alone.

We conducted a theoretical investigation into parallel genetic evolution and speciation from standing variation across the continuum from parallel to divergent natural selection. We primarily used individual-based simulations and include some simple analytical arguments to gain intuition. We compared results from simulations where adaptation proceeds simultaneously via the sorting of ancestral standing genetic variation and *de novo* mutation to simulations where adaptation proceeds via *de novo* mutation alone. Our results provide insight into the circumstances under which we should expect high vs. low genetic parallelism and also suggest that standing variation has substantial and unexpected implications for speciation that depend on the difference in the direction of natural selection between populations.

## Methods

We used computer simulations to investigate genetic parallelism and progress toward speciation (via ecologically-dependent postzygotic reproductive isolation) from standing variation across the continuum from parallel to divergent natural selection. Our simulations consider pairs of populations and multivariate phenotypes determined by multiple additive loci. In our simulations, a single ancestral population founds two identical populations that each adapt in their respective environments without gene flow (i.e., allopatry; see Fig. 1A). After adaptation, populations interbreed to form recombinant hybrids. This general colonization history—a single population splitting into two populations that adapt to their respective novel environments in allopatry—is modelled around the process of adaptation as it can occur in nature, for example in postglacial fishes (Bell and Foster 1994) and in birds or plants isolated within glacial refugia (e.g., Weir and Schluter 2004; Pettengill and Moeller 2012). In many of these cases, ecologically-dependent postzygotic isolation is thought to be essential for maintaining reproductive isolation (Nosil 2012). See Table 1 for descriptions of all parameters and values used in simulations (see **Data Accessibility** for information about accessing simulation code).

**Figure 1.**
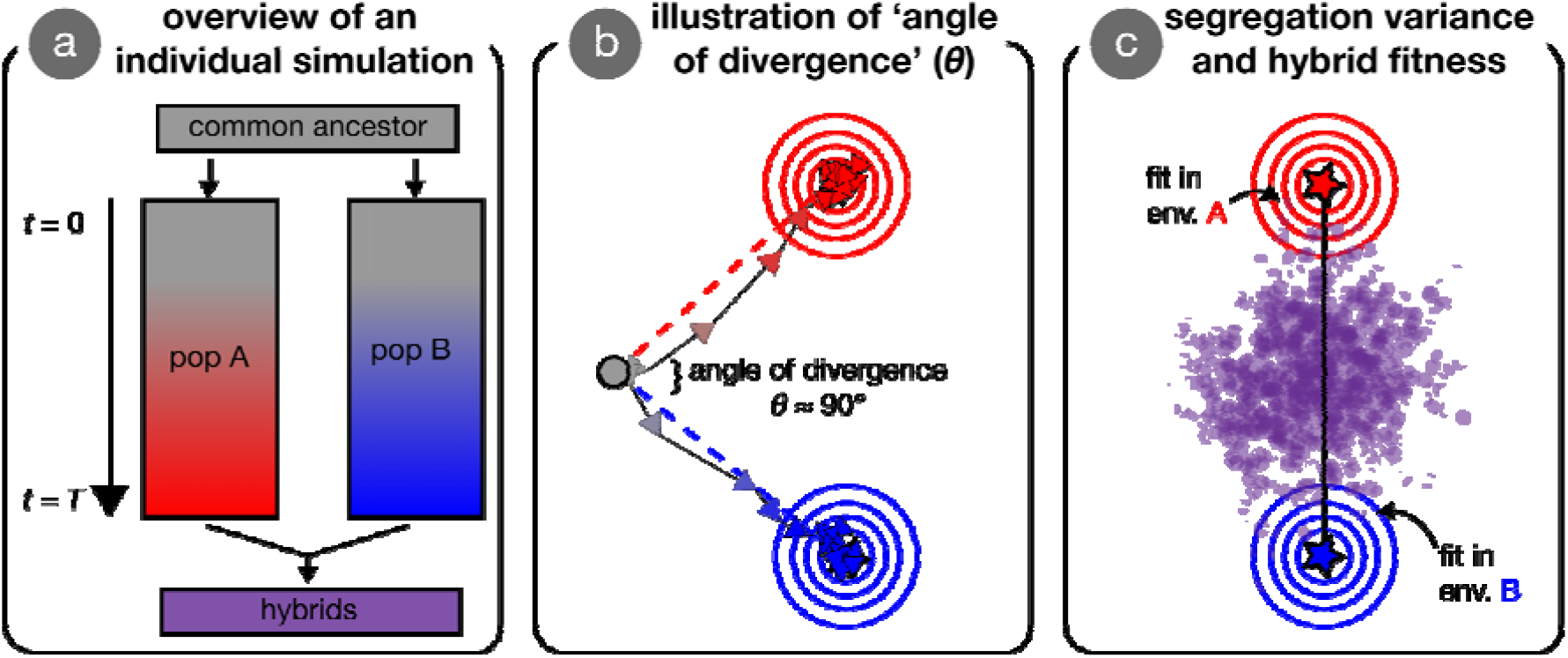
Visual overview of simulations and concepts. Panel (a) provides an overview of an individual simulation run. An ancestral population founds two initially-identical parental populations, that evolve independently for *T* generations in their respective environments. After *T* generations of adaptation, these parental populations interbreed to form hybrids. Panel (b) illustrates the process of adaptation in our simulations, wherein two populations (red and blue arrows connect the mean phenotype every 200 generations) independently adapt to specified optima (red and blue stars; behind arrows in [b] but visible in [c]). Concentric circles represent fitness contours around the two optima. The ancestor state is indicated by the grey dot, with the angle of divergence, *θ,* shown between the two axes of selection (red and blue dashed lines; angle shown is approximately 90°). Panel (c) illustrates the segregation variance in a group of hybrids. Individual hybrids (purple points) that are near an optimum have high fitness when measured in that environment. The black line is the line connecting parental optima—variance along this line can increase mean hybrid fitness whereas variance orthogonal to this line is necessarily deleterious.

**Table 1.**
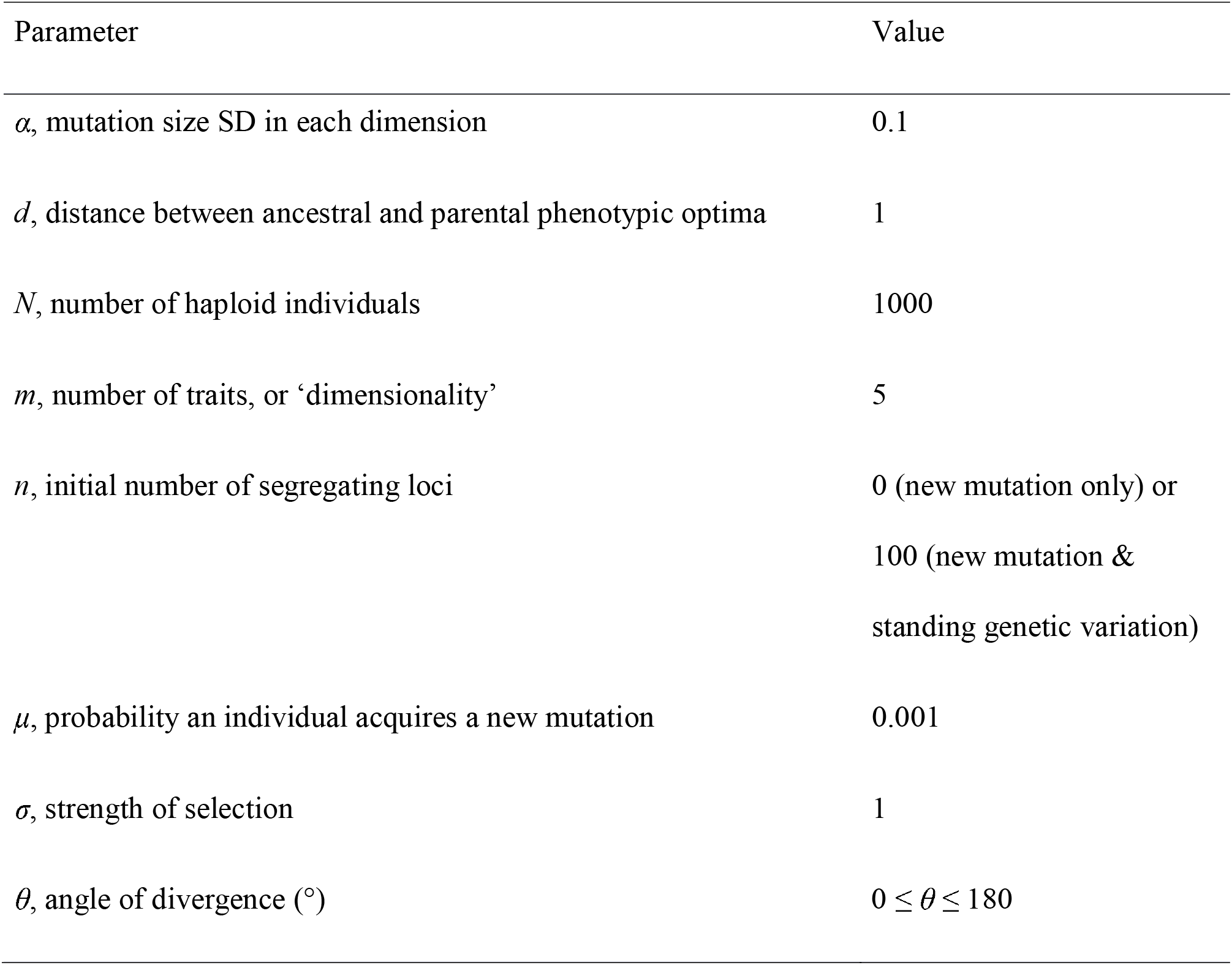
Description of parameters and parameter values in parental populations for simulations presented in the main text.

### Genotype to phenotype

The phenotype of a haploid individual is represented by an *m*-dimensional vector, ***z*** = [*z*_1_, *z*_2_,…, *z_m_*], with *m* being the number of uncorrelated ‘traits’ or phenotypic ‘dimensions’ (for further discussion of ‘dimensionality’, see Orr [2000] & Tenaillon [2014]). Each trait value, *z_i_*, is determined by the summed effects of alleles at all underlying loci (i.e., mutations act additively to determine the phenotype), which are initially fixed for alleles with an effect of 0 on all *m* traits. We present results from simulations with five phenotypic dimensions (*m* = 5) in the main text, and results for alternative parameter combinations are found in the supplementary figures (Figs. S1-S7).

### Life-cycle

We model a Wright-Fisher population (Fisher 1930; Wright 1931) with haploid selection. Fitness is a Gaussian function that depends on the Euclidean distance between an individual’s phenotype and the phenotypic optimum, ||***z*** − ***o***||, and the strength of selection, σ (e.g., Lande 1979):

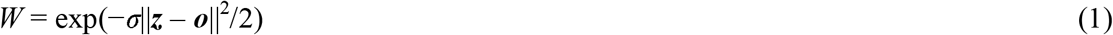

*N* haploid parents are then randomly sampled with replacement from a multinomial distribution with probabilities proportional to their fitness, *W*. Parents then randomly mate and produce two haploid offspring per pair, with free recombination between all loci. With probability *μ* an offspring gains a mutation; we assume an effectively infinite number of loci such that all mutations arise at a previously unmutated locus (‘infinite-sites’ *sensu* Kimura [1969]).

Mutational effects are drawn from a multivariate normal distribution (‘continuum-of-alleles’ *sensu* Kimura [1965]), with a mean of 0 and an SD of *α* in all *m* traits and no correlations among traits (i.e., universal pleiotropy). Our qualitative conclusions are robust to alternative assumptions about fitness functions (Fig. S8).

### Generating ancestral standing genetic variation

We initiate two parental populations with standing genetic variation from a common ancestor that adapt to novel phenotypic optima. To generate ancestral standing variation, we conducted burn-in simulations of a large ancestral population (*N*_anc_ = 10,000) under stabilizing selection (σ_anc_ = 0.01) at the origin (*o*_anc_ = [0, 0, − 0]) for 100,000 generations. All other parameters in the ancestor (e.g., mutation rate) were identical to those of the pair of parental populations that it founds. This parameter combination facilitates the accumulation of appreciable standing variation (see Fig. S9), but we note that our general conclusions also hold if the ancestor is under much stronger selection (σ_anc_ = 1) that puts it into the multivariate ‘House-of-Cards’ regime (Turelli 1985; see Fig. S10).

Ancestral populations reached mutation-selection-drift balance such that the rate of acquisition of new mutations was balanced by the rate of loss of mutations that arose in earlier generations (Fig. S9A). The mean frequency of derived alleles and the phenotypic (genotypic) variance was stable (Fig. S9B), as has been found in other models of phenotypes under stabilizing selection (e.g., Barton 1989). Segregating derived alleles were all at unique loci (by assumption) and were at low frequency in the ancestral population (see Fig. S9D for the site frequency spectrum). High derived allele frequencies and fixation are sometimes reached by drift when mutations have nearly-neutral selective coefficients and by positive selection when mutations compensate for deleterious alleles that have risen to high frequency by drift (Hartl and Taubes 1996; Orr 2005).

### Adaptation to a new environment

In simulations with standing genetic variation, a parental population was established by first randomly choosing *n* polymorphic loci in the ancestor (see Fig. S11 for effect of *n* on genetic parallelism and segregation variance). Each parental individual received the mutant (i.e., ‘derived’) allele at each of these *n* loci with a probability equal to the allele’s frequency in the ancestor. Loci fixed in the ancestral population were also fixed in the parental population. This rather artificial sampling procedure allowed us more control over the amount of standing genetic variation across simulations with different parameter values (Figs. S1-S7). Further control was achieved by making the second parental population initially identical to the first, so that each possessed the exact same collection of genotypes and there were therefore no founder effects. Populations adapted from only new (i.e., *de novo*) mutation when *n* = 0. Within each parameter combination, we began each replicate simulation from a unique realization of the ancestor (i.e., distinct burn-in). After initialization, parental populations adapted to their respective phenotypic optima without inter-population gene flow (Fig. 1B), and adaptation proceeded via natural selection on ancestral standing variation (if *n* > 0) and new mutation simultaneously.

Two properties of the new phenotypic optima are key. The first is the Euclidean distance between each optimum and the origin, *d* (assumed the same for both parental populations for simulations presented in main text). More distant optima yield a greater amount of genetic and phenotypic change. In the main text we set *d* = 1, which is equivalent to 10 times the SD of mutation effect size (*α*). The second key feature of the new optima is the angle of divergence, *θ*, separating vectors that originate at the origin and each pass through one of the parental optima (dashed lines in Fig. 1B). Angle is used to quantify the difference in the direction of selection from parallel (*θ* = 0°) to divergent (*θ* = 180°) and is explicitly invoked in most empirical metrics that quantify phenotypic parallelism (see Bolnick et al. 2018). The value of *θ* is what determines the mean phenotypic differences that evolve between parental populations in these simulations (because *d* is held constant).

We ended the adaptation phase of simulations after *T* generations, at which time all populations had reached their phenotypic optima (Fig. S12A) and mutation-selection-drift balance (Fig. S12B). An unavoidable and important effect of standing variation is that it quickens adaptation because populations do not have to ‘wait’ for beneficial alleles to arise (Barrett and Schluter 2008). In our model and others like it (e.g., Barton 2001 & Chevin et al. 2014), reproductive isolation evolves rapidly during the initial stages of adaptation. After populations reach their respective phenotypic optima, genetic divergence accumulates slowly at a rate proportional to the mutation rate (Barton 1989, 2001; Chevin et al. 2014). Therefore, our general conclusions reflect quasi-equilibrium conditions rather than transient states and are unaffected by standing variation’s influence on the speed of adaptation.

### Quantification of genetic parallelism and hybrid segregation variance & fitness

After the adaptation phase of simulations had ended, we calculated the proportion of alleles that fixed in both populations (i.e., genetic parallelism). We then we paired random individuals from the two parental populations to produce 100 recombinant haploid F_1_ hybrids. In the hybrids, we quantified the phenotypic segregation variance and the relative mean fitness compared to parents.

To quantify parallel genetic evolution between parental populations, we first determined the number of alleles that fixed in each population (Nfix_P1_ & Nfix_P2_) and the number of alleles that fixed in both populations (Nfix_Both_). We then calculated our metric of ‘genetic parallelism’ as:

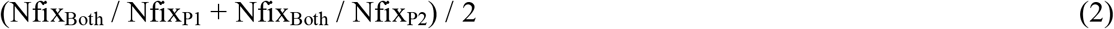

Values of 1 indicate complete genetic parallelism (i.e., all alleles that fixed were fixed in both populations) and values of 0 indicate complete genetic non-parallelism (i.e., no allele was fixed in both populations). We use this metric because of its ease of interpretation and note that it is highly correlated with other metrics of genetic divergence between populations (e.g., F_ST_; Fig. S13).

After forming hybrids we quantified their phenotypic variation—the net segregation variance (Wright 1968; Slatkin and Lande 1994)—calculated here as the mean phenotypic variance across all *m* traits. We present analyses of individual axes of variance where relevant. Higher segregation variance results when parents are differentiated by a greater number of alternative alleles (holding effect size constant) or alleles of individually-larger effect (holding number of alleles constant) (Castle 1921; Slatkin and Lande 1994; Chevin et al. 2014). Segregation variance captures the phenotypic consequences of hybridization and has a direct impact on fitness whereas genetic (non)parallelism is only indirectly related to fitness. Phenotypic variance in parental populations before hybridization is near zero and does not differ between populations founded with vs. without standing variation nor does it depend on the initial distance to the optimum (*d*; Fig. S12C).

An individual hybrid’s fitness in a given parental environment was calculated from its phenotype in the same manner as the fitness of parental populations (Fig. 1C). We determined the fitness (equation 1) of each hybrid in both parental environments and recorded its fitness as the larger of the two values. This can be imagined as, for example, giving the hybrid a choice of alternative host-plants (Drès and Mallet 2002), where the choice is always for the host to which it is better adapted. Our fitness metric reflects what is traditionally recognized as ‘extrinsic’ postzygotic isolation (Coyne and Orr 2004), and explicitly considers environment-specific epistasis for fitness (Bateson 1909; Dobzhansky 1937; Muller 1942; Chevin et al. 2014; Fraïsse et al. 2016; see also Arnegard et al. 2014; Schumer et al. 2014; and Ono et al. 2017 for discussion of environment-specific hybrid incompatibilities). Because hybrids are recombinant, hybrid fitness reflects both the effects of displacement of the mean phenotype from the optimum (the ‘lag’ load) and what in diploids is known as ‘F_2_ hybrid breakdown’ (Burton et al. 2006). We report hybrid fitness relative to the parents for each individual simulation, calculated as: [mean fitness of hybrids] / [mean fitness of parents].

## Results

### Genetic parallelism and phenotypic segregation variance along the continuum from parallel to divergent selection

We first investigate how genetic parallelism between two populations is associated with the angle of divergence (*θ*) when adaptation is from standing variation (dark green line and points). Genetic parallelism is highest under completely parallel natural selection (*θ* = 0°) and rapidly decreases toward its minimum value as θ increases (Fig. 2A; see black line for visual comparison of deviation from linearity). This rapid decrease in genetic parallelism also occurs when the phenotypic distance *between* optima is used as the independent variable, although we note that non-linearity is only appreciable in higher dimensions (see Fig. S14). There is considerable variation in genetic parallelism even when populations adapt to identical environments, which presumably results from stochastic processes in each run. For example, alleles are lost due to drift, populations fix weakly deleterious alleles or different *de novo* mutations, and populations fix alternative alleles from the standing variation early in the simulations which affects the selection coefficients of all other alleles in later generations (Chevin and Hospital 2008). Genetic parallelism never decreases to zero even under completely divergent selection (*θ* = 180°), indicating that populations fix some deleterious alleles during the course of adaptation (although genetic parallelism is nearer to zero in higher dimensions; Fig. S1-3). Our conclusion that genetic parallelism rapidly decreases with θ are generally robust to variation in population size and selection strength, except for when small populations are under weak selection (Fig. S1), likely due to an overwhelming effect of drift (see Fig. S15 for divergence between populations due to drift alone at various population sizes).

**Figure 2.**
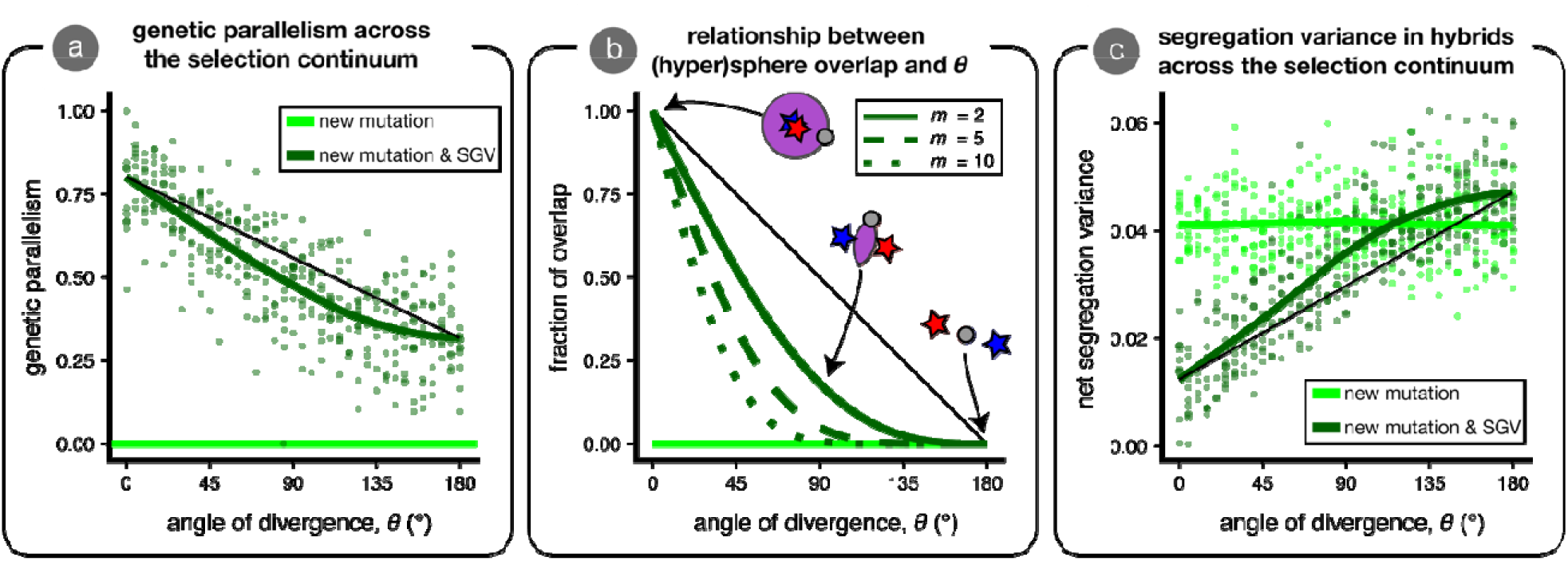
Genetic parallelism and phenotypic segregation variance across the continuum from parallel to divergent natural selection. We conducted 10 replicate simulations for optima separated by different angles (*θ* = 0° to 180°; *d* = 1). Parental populations adapted from either new mutation only (n = 0; light green) or from a combination of new mutation and standing genetic variation (SGV) (n = 100; dark green). Panel (a) shows the proportion of alleles that fixed in both populations (genetic parallelism; equation 2). Green lines are loess fits. The thin black line connects the fit at *θ* = 0° to the fit at *θ* = 180° and is shown to facilitate visualization of the nonlinearity. Panel (b) is an analytical result that depicts the relationship between *θ* and the fraction of overlap between two (hyper)spheres for three different dimensionalities (*m*) (see equation A1). In the inset cartoon, mutations that bring the phenotype into the red and blue regions are initially beneficial only in the ‘red’ or ‘blue’ environments, while mutations that bring the phenotype into the purple region are beneficial in both environments. The horizontal light green line is set at 0 where there is no overlap. Panel (c) shows the net phenotypic segregation variance in hybrids.

Genetic parallelism decreases with *θ* because the fraction of alleles that are beneficial in each of the two populations declines as *θ* increases. For a given population, beneficial alleles bring populations closer to the middle of a hypersphere centred at the phenotypic optimum (Fisher 1930; see cartoon inset of Fig. 2B). Considering two populations, each with their own hypersphere, a given allele is beneficial in both—and thus could fix in parallel via positive natural selection—if it brings a population’s phenotype into the region where the two hyperspheres overlap (purple region in Fig. 2B inset). The size of this *region of overlap* decreases rapidly with *θ* (Fig. 2B; see Appendix for mathematical details), and therefore so does the fraction of alleles present as standing variation that are beneficial in both populations. The rate of decrease of overlap is faster with greater dimensionality (compare solid line to dashed lines in Fig. 2B; see Figs. S1-S3 for confirmation with simulations in higher and lower dimensions) but—perhaps surprisingly—does not depend on the distance to the optima (*d*; if *d*_1_ = *d*_2_ = *d*) and is not expected to change over the course of an ‘adaptive walk’ (*sensu* Orr [1998]; see Appendix and Fig. A1 for detailed explanation).

It is possible to imagine an alternative case where *θ* is held constant, but populations differ in the distance to their respective optima (i.e., different vector ‘lengths’ rather than ‘angles’ *sensu* Bolnick et al. [2018]). Even if selection acts in the exact same direction on two populations (*θ* = 0°; *m* = 5), if the phenotypic optimum of population 2 is twice as far as that of optimum of population 1 from the ancestral phenotype, less than 5% of the alleles beneficial to population 2 are also beneficial to population 1 (see Fig. S16). The region of overlap contains only small effect alleles, however, which tend to be the sort of alleles that are present in the standing variation. Therefore, the extent of genetic parallelism from standing variation might be higher than what might be expected in a case where adaptation is from *de novo* mutation alone.

The changes in segregation variance generally mirror patterns of genetic parallelism (Fig. 2C). With standing variation, segregation variance is low under parallel selection and rapidly increases with *θ.* Although Chevin et al. (2014) found that segregation variance (proportional to their ‘variance load’) does not depend on *θ,* this is because their model did not permit genetic parallelism. When there is no standing variation, segregation variance is not affected by the angle of divergence (light green line and points in Fig. 2C; linear model *P* > 0.9 [as a check]), in agreement with the findings of Chevin et al. (2014; their Fig. 2). At large angles, net segregation variance is greater when populations adapt from standing variation than when they adapt from new mutation alone, and the magnitude of this difference increases with dimensionality (see Fig. S17). We explore the reasons for this below.

### Effect of standing variation on hybrid fitness across the continuum from parallel to divergent selection

In this section, we evaluate the effect of standing variation on hybrid fitness across the continuum from parallel to divergent natural selection. The most readily observable pattern is that the mean relative fitness of hybrids is lower under divergent selection than under parallel selection regardless of whether adaptation proceeds with standing variation (Fig. 3A). This pattern occurs mainly because the hybrid mean phenotype is increasingly distant from either parental optimum as *θ* increases. In Fig. 3A, we plot the fitness of the hybrid mean phenotype (the ‘lag’ load) as a thin black line.

**Figure 3.**
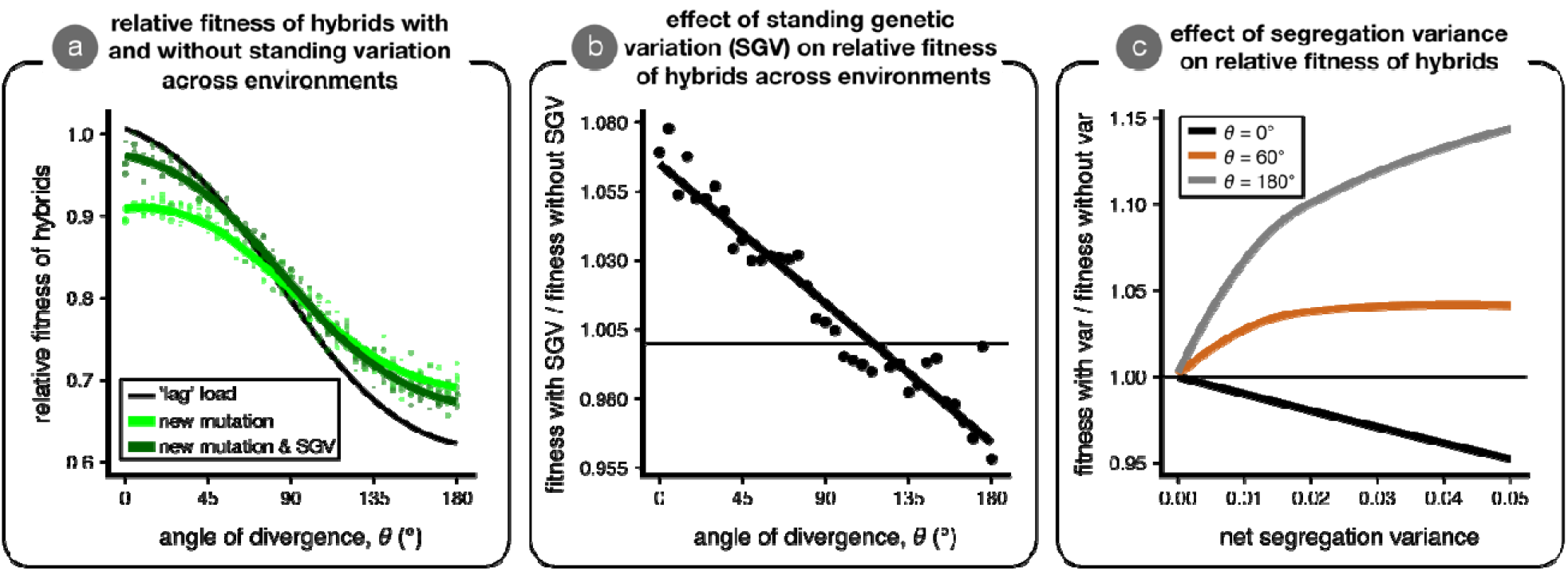
The effect of standing variation on mean hybrid fitness across the continuum from parallel to divergent natural selection. Panel (a) shows the mean relative fitness of hybrids—as compared to parents—across environments in simulations initiated without (*n* = 0; light green) and with (*n* = 100; dark green) ancestral standing genetic variation. The thin black line represents the mean relative fitness of hybrids due only to the deviation of the observed mean phenotype from an optimum (‘lag’ load), and is close to 1 when the hybrid mean phenotype is on the optimum. Panel (b) shows the effect of standing variation on mean relative hybrid fitness (the ratio of values for dark / light green lines in panel [a]); the horizontal line indicates no effect of standing variation on relative mean hybrid fitness. Panel (c) is an analytical result that illustrates the relationship between segregation variance and mean hybrid fitness for three angles of divergence (black, *θ* = 0°; brown, *θ* = 60°; grey, *θ* = 180°) when the hybrid phenotype is multivariate normal with a mean exactly in between the two parental optima and equal variance in all phenotypic dimensions (no covariance). Hybrid fitness is plotted for each angle relative to the case of no variance; the horizontal line indicates when segregation variance has no effect on hybrid fitness.

Compared to when adaptation is from new mutation, adaptation from standing variation improves mean hybrid fitness when parental populations adapt to similar optima but can reduce hybrid fitness when parents undergo divergent adaptation (Fig. 3B). This pattern is caused by environment-specific effects of segregation variance on mean hybrid fitness (Fig. 3C, Fig. 4). When the hybrid phenotype distribution is centred at the phenotypic optimum, as it is under parallel selection (*θ* = 0°), segregation variance is universally deleterious. When parental populations adapt to identical optima from only new mutation, hybrids vary considerably around the parental optimum and thus have relatively low fitness. When populations have access to a common pool of standing variation, parallel genetic evolution leads to lower segregation variance around the optimum and therefore higher mean fitness under parallel selection (Fig. 4A; see Fig. S18 for similar results but for *maximum* hybrid fitness instead of mean).

**Figure 4.**
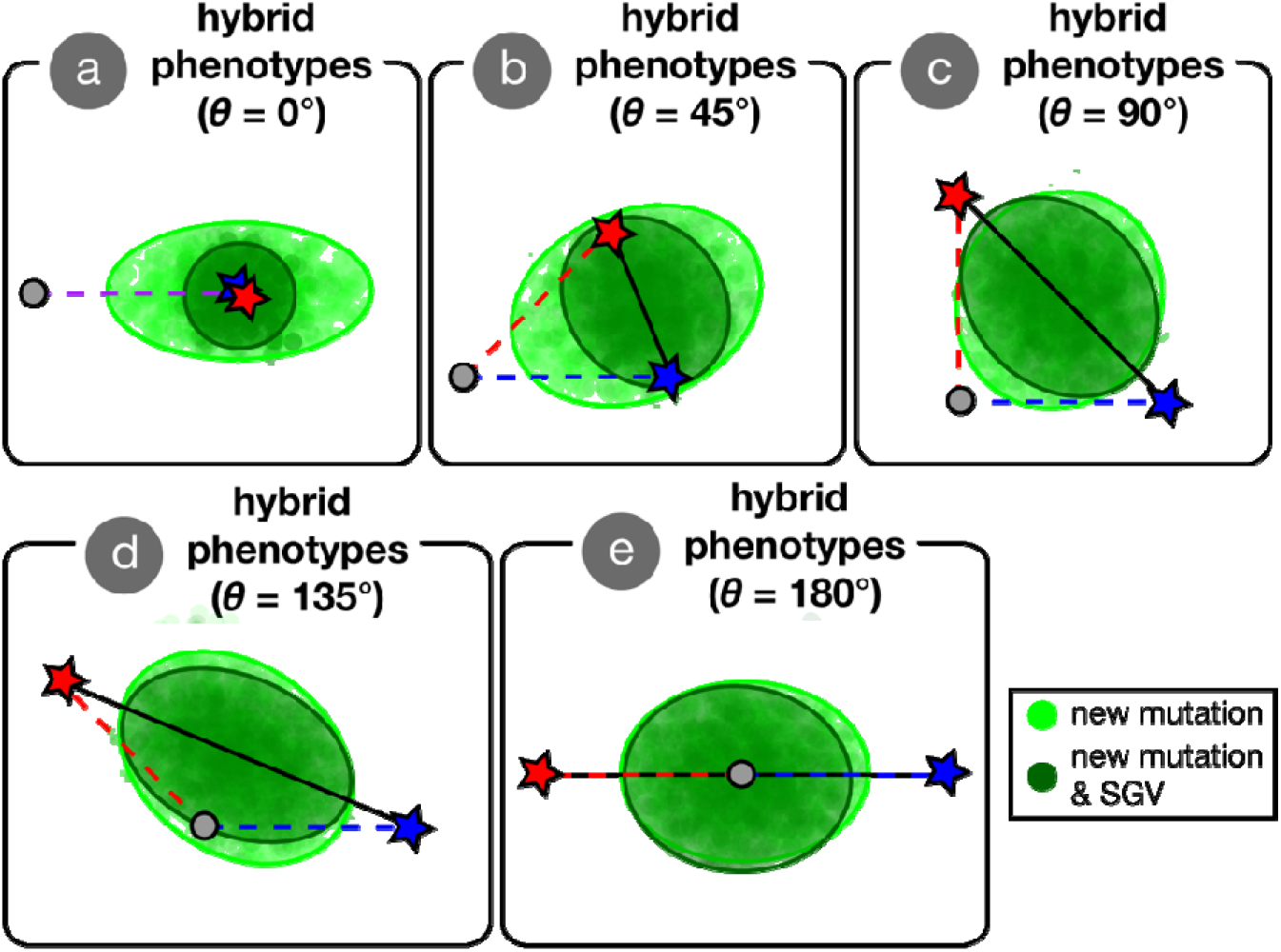
The effect of standing variation on the distribution of hybrid phenotypes. We plot hybrid phenotypes (points) and 95 % confidence ellipses for five angles of divergence (*θ*) evenly spaced along the continuum of (a) completely parallel (0°) to completely (e) divergent (180°) for simulations where populations adapted from only new mutation (light green) or both new mutation and standing genetic variation (dark green; 1000 hybrids each from ten replicate simulations each producing 100 hybrids). Parental optima are depicted as stars and the origin (ancestral optimum) is shown as a grey dot. The axes of selection connect the origin and optima (dashed red and blue lines) and we also show the axis connecting parental optima as a black line. We show the first two (of five) phenotypic dimensions, which are the only dimensions in which the optima might differ.

At large angles of divergence, adaptation from standing variation reduces hybrid fitness compared to when adaptation is from only new mutation—a result we did not anticipate. The general reason for this is that hybrids (but not the parents) increasingly fall in a ‘fitness valley’ between parental optima as the angle of divergence increases, and when in this valley some segregation variance is beneficial for mean hybrid fitness (see Fig. 3C). This result is robust to variation in parameter values (see Figs. S4-S6), except for when selection is very weak in small populations. We plot the distribution of hybrid phenotypes resulting from representative simulations across the continuum from parallel to divergent selection in Figure 4. From these figures, it is clear that there are appreciable differences in patterns of phenotypic variation in hybrids when their parents adapt with standing variation vs. when adaptation is from new mutation alone. Only phenotypic variation along the axis connecting parental optima (black line connecting stars in Fig. 4) is beneficial, whereas variation along orthogonal axes is deleterious. When *θ* = 180°, hybrids have reduced variation along the axis connecting parental optima and slightly more variation along all other axes. Thus, maladaptive segregation variance reduces hybrid fitness under large angles of divergence.

Why does adaptation from standing variation alter patterns of phenotypic segregation variance in hybrids? As discussed above, adaptation from standing genetic variation reduces segregation variance under parallel selection because parents fix the same alleles that therefore do not segregate in hybrids. Populations adapting from standing variation also fix a greater number of smaller effect alleles than simulations initiated without standing variation (Fig. 5A & B), which contributes to the reduction in hybrid segregation variance along the mean axis of selection in parents (average of dashed and blue lines in Fig. 4). Fixation of smaller-effect alleles likely occurs under adaptation from standing variation because stabilizing selection removes large-effect alleles from the standing variation (Fig. S9) and because weakly beneficial alleles have a higher probability of fixation when present in standing variation compared to if they arose *de novo* (Orr and Betancourt 2001; Hermisson and Pennings 2005; Matuszewski et al. 2015).

**Figure 5.**
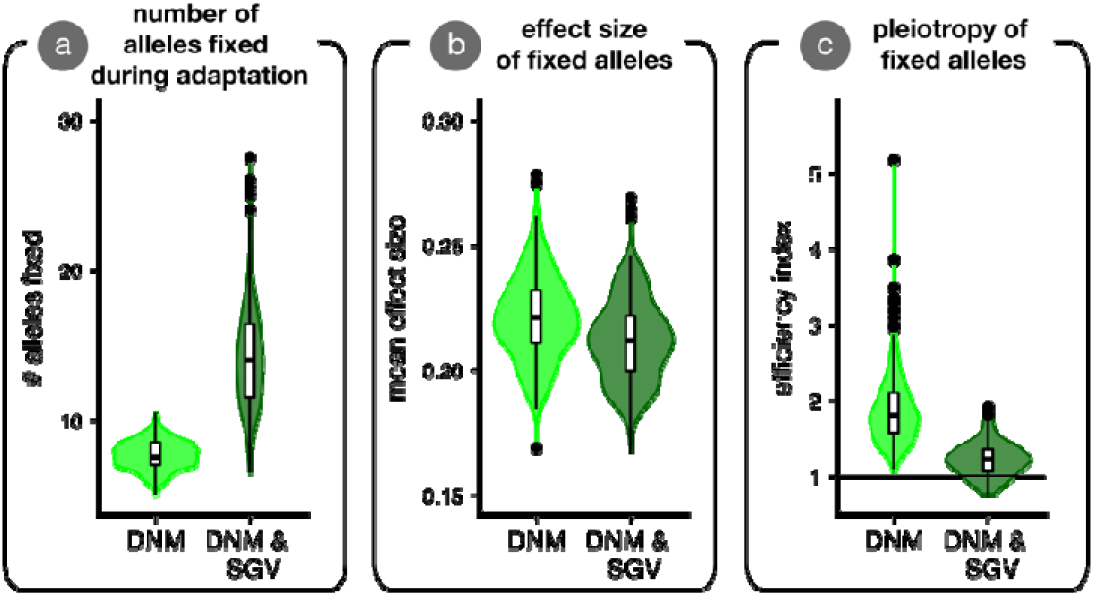
Properties of alleles fixed during adaptation. We show results from simulations where parental populations adapted from only *de novo* mutation (light green; DNM) vs. adaptation from standing variation and new mutation (dark green; DNM & SGV). Each replicate simulation contributed one data point to the plot. Panel (a) shows the average number of alleles fixed during adaptation. Panel (b) shows the average effect size (Euclidean length of mutation vector) of alleles fixed during adaptation. Panel (c) shows the allele ‘efficiency index’, which plots the ratio of a fixed mutations’ effect size in the direction of selection vs. orthogonal directions. Values of 1 are equally balanced in these directions, and mutations are more ‘efficient’ (i.e., they point more directly at the optimum) as this index increases. Statistical tests confirm all differences as highly significant (not shown).

This latter effect seemed to allow alleles with more deleterious pleiotropic effects to fix during adaptation from standing variation than when adaptation was from new mutation alone (Fig. 5C). We quantified pleiotropy by taking the ratio of the mean effect size of fixed alleles along the axis of selection in parents (red or blue dashed lines in Fig 4) vs. the mean effect size across all orthogonal axes, termed the ‘efficiency index’. Values of 1 (horizontal line in Fig. 5C) imply that an allele had an equivalent effect along the axis of selection as on any orthogonal axis. Increasingly positive values reflect alleles that take a population to the optimum more ‘efficiently’ (i.e., directly along the dashed blue or red line in Fig. 4). Together, these results indicate that adaptive walks from standing variation in our simulations involved more—slightly smaller—steps and are more ‘meandering’ than adaptive walks from new mutation alone, which use fewer, slightly larger, and more direct steps (but see Ralph and Coop 2015). These differences in the properties of alleles fixed in simulations initiated with vs. without standing variation contribute to the patterns of phenotypic segregation variance that influence mean hybrid fitness.

## Discussion

In this study we investigated parallel genetic evolution and progress toward speciation under adaptation from standing variation. Using a combination of individual-based simulations and analytical models, we characterized how the extent of genetic parallelism from standing variation changes with the angle of divergence. We then illustrated how adaptation from standing variation affects hybrid fitness across the continuum from parallel to divergent natural selection, compared to when adaptation is from only new mutation. Here, we highlight our key findings, predictions for empirical systems, and discuss suggestions for future work.

### Key predictions and possible tests

The first principal finding of our study is that the degree of genetic parallelism rapidly declines as the angle of divergence increases from parallel toward divergent, especially when a large number of traits affect fitness. Practically, this means that the extent of genetic parallelism also declines quickly with phenotypic divergence. It is possible to test this prediction in natural or experimental populations using techniques such as ‘Phenotypic Change Vector Analysis (PCVA)’, which estimates important parameters such as the angle between the vectors and/or the difference in their magnitudes (Bolnick et al. 2018). For example, hundreds of populations of threespine stickleback (*Gasterosteus aculeatus*) have adapted to freshwater lakes and streams from a shared marine ancestor (Jones et al. 2012). Appreciable phenotypic differences exist among freshwater-adapted populations (Bell and Foster 1994), and if we assume that marine populations represent the ‘ancestral’ state that founded ‘derived’ freshwater populations (see Morris et al. 2018 for analysis of regional variation of the ‘contemporary ancestor’), parameter estimates of marine-freshwater divergence from PCVA will give some indication about what the genetic parallelism underlying marine-freshwater divergence is expected to be. Other cases with repeated instances of easily-quantified phenotypic divergence (see Oke et al. 2017; Stuart et al. 2017) are also amenable to this approach. Given that phenotypic and genetic parallelism are not linearly related (Fig. S14), we suggest that analytical predictions about the extent of genetic parallelism ought to be considered when generating predictions for empirical systems.

Our second principal finding is that—relative to when adaptation is from only *de novo* mutation—adaptation from standing genetic variation improves the mean fitness of hybrids under parallel natural selection, has little effect at intermediate angles of divergence, and reduces mean hybrid fitness under completely divergent selection. Practically, this indicates that adaptation from standing variation works against ‘mutation-order’ speciation and facilitates ‘ecological’ speciation (Schluter 2009; Schluter and Conte 2009). This hypothesis could be tested most readily in experimental systems where the amount of ancestral standing variation can be easily manipulated, and where interpopulation hybrids can easily be generated to have their fitness measured in parental environments.

### Alternative sources of standing variation

Our model addresses the case of adaptation from a pool of standing genetic variation at mutation-selection-drift balance. This framework does not address cases of adaptation where standing variation is generated from other sources. For example, in threespine stickleback, the marine ancestral form is thought to maintain standing variation for freshwater-adapted alleles in a balance between migration of alleles from freshwater populations and negative selection in the sea (the ‘transporter’ hypothesis; Schluter and Conte 2009; Nelson and Cresko 2018). In this case, the pool of standing variation is enriched for alleles that have already swept to fixation in freshwater populations—that is, they are ‘pre-tested’ by selection—which might occur in linked blocks of freshwater-adapted alleles in the sea. Scenarios such as this are especially likely to cause genetic parallelism, more than is predicted from sort of standing variation modeled here (Schluter and Conte 2009). This might help to explain why genetic parallelism is so high between independently-evolved freshwater populations of stickleback that have independently adapted from a common ancestral phenotype (Jones et al. 2012). The extent to which adaptation from standing variation proceeds via the sorting of ‘naïve’ vs ‘pre-tested’ alleles is unresolved.

### Possible extensions

Our study represents a step towards characterizing changes in genetic parallelism and progress toward speciation in pairs of populations experiencing a variety of differences in the direction of natural selection between them. Many of our assumptions—for example a lack of recurrent *de novo* mutation or gene flow—reduce the extent of genetic parallelism (Nosil and Flaxman 2011; Anderson and Harmon 2014; Ralph and Coop 2015). In addition, we considered only haploid selection, no dominance, universal pleiotropy, and strict additivity of allelic effects on phenotypes (i.e., no epistasis). We also assumed that the sole fitness optima available to hybrids are those that the two parents are adapted to (see Rieseberg et al. 1999). Our analytical results consider only the fraction of mutations that are mutually beneficial, ignoring differences in the probability that particular mutations arise and fix. Extending our approach to integrate the distribution of fitness effects of new mutations (Eyre-Walker and Keightley 2007), the correlation of selection coefficients across environments (Kassen 2014; Martin and Lenormand 2015), and existing theory on the probability of genetic parallelism from standing variation (MacPherson and Nuismer 2017) will be valuable.

We also note that the only reproductive isolating barrier we considered was environment-specific post-zygotic isolation. Post-zygotic isolation can also be environment-independent, and such ‘intrinsic’ isolating barriers are correlated with genetic divergence between populations (Orr 1995; Matute et al. 2010; Moyle and Nakazato 2010; Wang et al. 2015). Chevin et al. (2014) quantified the strength of intrinsic postzygotic isolation using a metric of ‘variance load’, which is proportional to our metric of net segregation variance. Therefore, our measure of segregation variance might be interpreted as being proportional to the strength of intrinsic barriers. We also did not consider pre-zygotic barriers such as assortative mating (Gavrilets 2004), which are also important for maintaining reproductive isolation. Accordingly, our results might be most relevant for empirical systems where ecology-based postzygotic isolation has a primary role in the origin of species.

### Concluding remarks

In this study we characterized patterns of genetic parallelism and progress toward speciation from standing variation in pairs of populations with quantitative differences in the direction of selection between them. Our findings generate new hypotheses for empirical studies on genetic parallelism and speciation. As evolutionary biologists develop increasingly powerful tools for detecting parallel genetic adaptation in nature, it will be important to keep in mind that genetic parallelism may be less common than we might intuit from patterns of selection and phenotypic similarity. We have also shown that adaptation from standing variation is expected to weaken the strength of isolating barriers that evolve between populations subject to parallel natural selection. By contrast, our simulations indicate that adaptation from standing variation can actually facilitate the process of speciation via divergent natural selection (i.e., ‘ecological’ speciation), suggesting that adaptation from standing variation might have a role in adaptive radiation beyond simply making it progress more quickly.

## Acknowledgements

Discussions with S. Otto motivated and refined the approach used in this study. Feedback from S. Arnold, M. Chapuisat, L. Chavarie, R. Holzman, A. MacPherson, S. Otto, L. Rieseberg, J. Rolland, M. Urquhart-Cronish, R. Yamaguchi and three anonymous reviewers improved the manuscript and/or reduced the demand for electricity by our simulations. K.A.T. was funded by The University of British Columbia, the Natural Sciences and Engineering Research Council of Canada (NSERC) and the Izaak Walton Killam Memorial Fund for Advanced Studies. M.M.O. was funded by The University of British Columbia. D.S. was funded by the Canada Foundation for Innovation, Genome BC, and NSERC.

## Author contributions

K.A.T. and D.S. developed the original ideas upon which the paper is based. K.A.T. wrote the first draft of the manuscript with input from M.M.O. and D.S., and all authors contributed to subsequent revisions. M.M.O. wrote the simulations and supplied analytical derivations with input from K.A.T. The simulations were performed by K.A.T. with input from M.M.O., and K.A.T. processed, plotted, and analyzed the data.

## Data accessibility

Python (version 3.6.4) scripts and resulting data, R (version 3.4.1) scripts to process and plot the simulated data, and a Mathematica notebook (version 9; and PDF copy) to derive analytical results will be archived on Dryad. For now, these are hosted on GitHub (https://github.com/Ken-A-Thompson/SVS).

## Supporting information for

### Patterns of speciation and parallel genetic adaptation from standing variation

### Appendix

Here we outline an explanation for why genetic parallelism decreases rapidly with the angle of divergence, *θ* (Fig. 2A, and distance between optima (Fig. S15B). Our explanation focuses on the extent of phenotypic space wherein mutations improve the fitness of both adapting populations in their respective environments. At the time of founding both adapting populations have the same mean phenotype, which is the mean ancestral phenotype. Mutations that move this ancestral mean phenotype into the region that leads to higher fitness in both parental environments are thus beneficial in both populations. The region of phenotypic space that has higher fitness than the mean phenotype in one environment is a hypersphere (of dimension m), centred on the optimum with a radius equal to the distance between the mean phenotype and the optimum, *d.* A similar hypersphere characterizes the phenotypic space that has higher fitness than the mean phenotype in the other parental environment. The region that is mutually beneficial is then the intersection of two hyperspheres, which is the union of two hyperspherical caps.

Fortunately, the volume of a hyperspherical cap is known for any dimension, *m* (Li 2011). It depends on the dimensionality (*m*), the radii of the two hyperspheres (*d*), and the distance between their centers (δ. In our case the distance between the two centres is *δ* = 2*d * sin*(*δ*/2). The amount of phenotypic space that is beneficial in a given environment is simply the volume of one of the hyperspheres. Thus, dividing the volume of the mutually-beneficial space (the union of the hyperspherical caps) by the volume of the space beneficial in a given environment (one of the hyperspheres) gives the fraction of beneficial mutations which are mutually beneficial. Using the formula given by Li (2011; their eqn 3) for the volume of a hyperspherical cap created by the intersection of two *m*-dimensional hyperspheres with radii *d* whose centres are distance *δ* = 2*d\** sin(*θ*/2) apart, the fraction of beneficial mutations that are expected to be beneficial in both is:

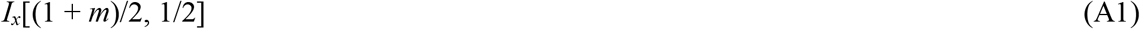

where *I_x_*[*a*, *b*] is the regularized incomplete beta function (Equation 6.6.2 in Abramowitz and Stegun [1972]) and here *x* = Cos(*ö*/2)^2^. Eq. A1 depends on only *m* and *θ*, that is the solution is independent of the distance from the ancestor to the new optima, *d*. We refer to Eq. A1 as the fraction of overlap in the main text, but note that this is only true when *d*_1_ = *d*_2_ (the formula is more complex when *d*_1_ ≠ *d*_2_, but can easily be used, e.g., Fig. S16B). The incomplete regularized beta function arises from integrating sin^*m*^(*$*) over *θ* (Li 2011).

The solution of Eq. A1 exhibits a rapid decrease with *θ* for all values of *m* > 0, and the decrease is faster for greater values of *m* (Fig. 2B). Thus, if standing genetic variation was uniformly distributed throughout the beneficial hyperspheres, the percent of segregating beneficial mutations that were beneficial in both parental populations, and thus expected to potentially fix in both, would decrease faster-than-linearly with the angle of divergence.

The above analysis considers only the very onset of adaptation, when the two parental populations have the same mean phenotypes, such that the fraction of phenotypic space that is beneficial in one population that is also beneficial in the other population (call this *X*) is equivalent to the fraction of possible beneficial mutations (if uniformly distributed across the hyperspheres) that are beneficial in both populations (call this *Y*). As adaptation proceeds the mean phenotypes of the parental populations depart from one another and *X* therefore no longer equals *Y*. This is because mutations are vectors that move a phenotype in a particular direction, and thus a mutually beneficial point in phenotypic space is only guaranteed to be a mutually beneficial mutation if both populations have the same mean phenotype.

To account for the inequality between phenotypic space (*X*) and mutational vectors (*Y*) during adaptation we must shift the mean phenotypes so that they are at the same point in phenotypic space and move their optima by an identical translation (see Fig. A1). We then have *X=Y*. One way to imagine this is to keep the mean phenotypes in place at the mean ancestral phenotype (the origin) and consider adaptation as the movement of the optima closer to the mean phenotypes. From this perspective, adaptation’s effect is a shrinking of the radii of the hyperspheres (at roughly equivalent rates in the two populations if adaptation proceeds relatively deterministically). Thus, because the fraction of overlap (Eq. A1) does not depend on the radii of the hyperspheres, the fraction of overlap is expected to remain constant throughout adaptation.

In reality and in our simulations, standing genetic variation is not uniformly distributed, the probability of fixation varies across the region of overlap, and adaptation uses up some of the standing variation so that the distribution of standing variation changes with time. Taking the first two complications into account would require weighted averages across the space contained in the hyperspherical caps, which is beyond the scope of our study. The third complication is yet more involved and would require an analysis of how standing genetic variation is used as adaptation proceeds (i.e., how the distribution of segregating effects and allele frequencies shift as alleles fix). Such a calculation is also beyond the scope of this article. Despite these complications, it seems as though the simple analysis above qualitatively captures the essence of why genetic parallelism decreases rapidly with the angle of divergence.

**Fig. A1.**
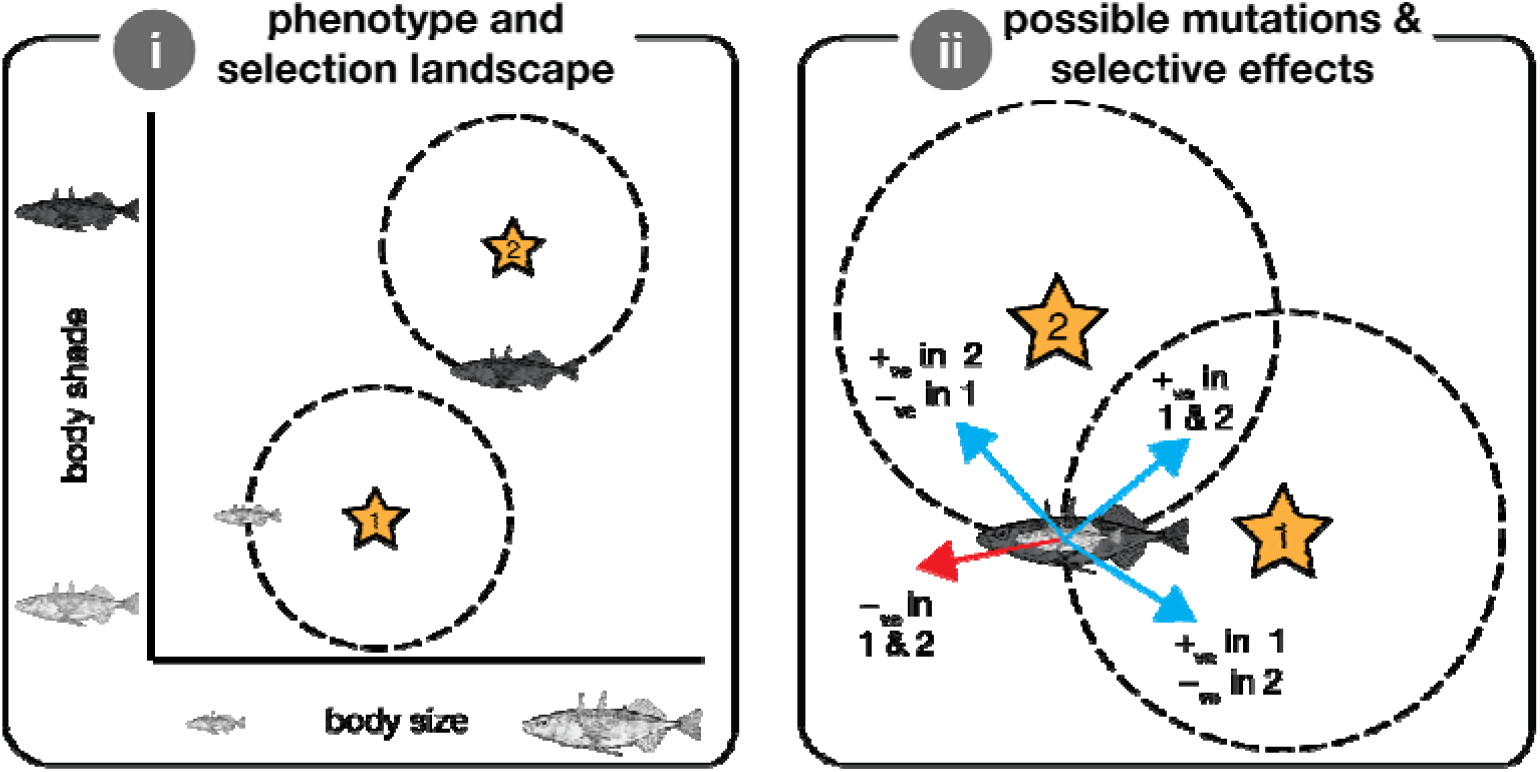
Cartoon illustration of why divergence among populations does not affect whether an allele is beneficial in both of them. Panel (i) depicts the phenotype landscape and selection landscape. Variation in the horizontal dimension reflects phenotypic variation in body size, and the vertical dimension reflects variation in body shade. We depict two ‘populations’ with differences in body size and shade (small & light; big & dark). The stars reflect local optima after a hypothetical environmental shift—selection favours adaptation toward a larger body size in population 1 and selection for darker body shade in population 2. If we illustrate the circle of beneficial mutational space (dashed circles) with respect to the current phenotypic position they do not overlap. Panel (ii) illustrates the selection landscape as it is ‘experienced’ by each population. An allele that slightly increases body size and darkens the body shade from the current phenotype (the position of the fish cartoons) is beneficial (blue) in both of populations. Some alleles are beneficial in only one population, and others are deleterious in both (red). Thus, even though the spheres do not overlap in (i) it is not the case that they populations will undergo non-parallel genetic evolution.

### Supplementary figures

**Figure S1.**
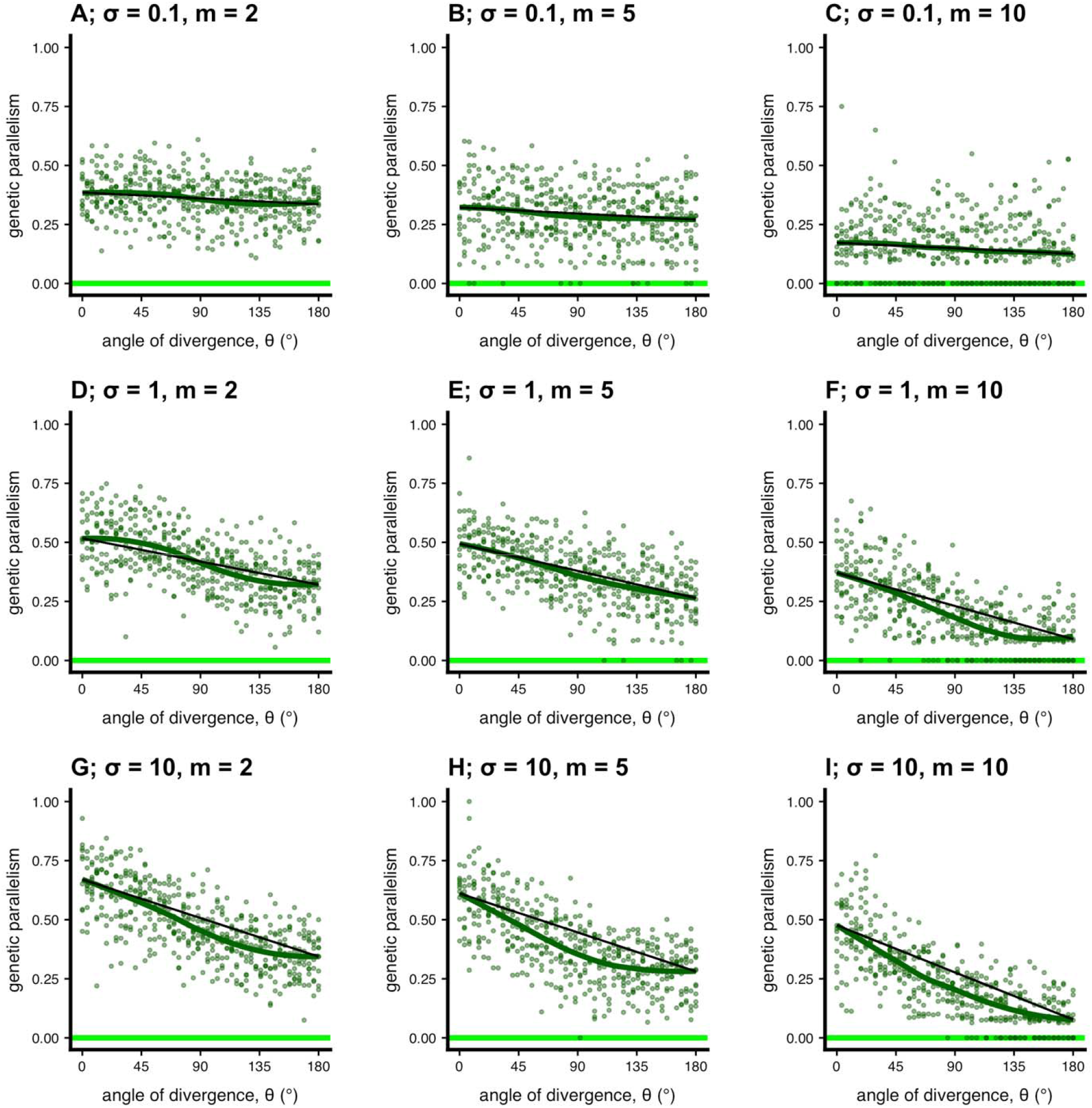
Genetic parallelism across the continuum of parallel to divergent natural selection (pop size = 100). This figure presents simulations similar to Fig. 2A in the main text but with varying parameter values (selection [*σ*] and dimensionality [*m*]). We ran these particular simulations for *T* = 5000 generations. All other parameters as in main text.

**Figure S2.**
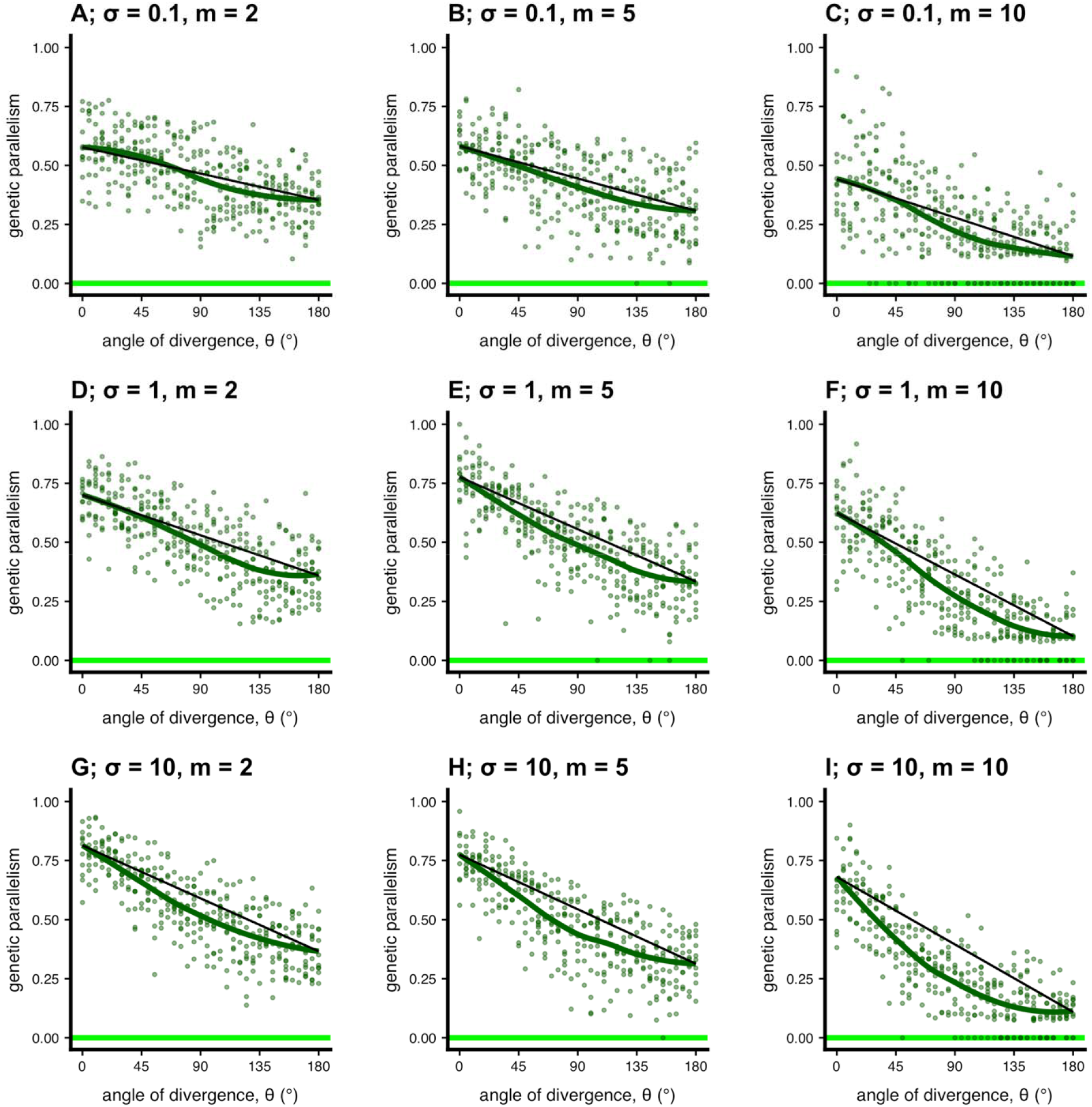
Genetic parallelism across the continuum of parallel to divergent natural selection (pop size = 1000). This figure presents simulations similar to Fig. 2A in the main text but with varying parameter values (selection [*σ*] and dimensionality [*m*]). We ran these particular simulations for *T* = 2000 generations. All other parameters as in main text.

**Figure S3.**
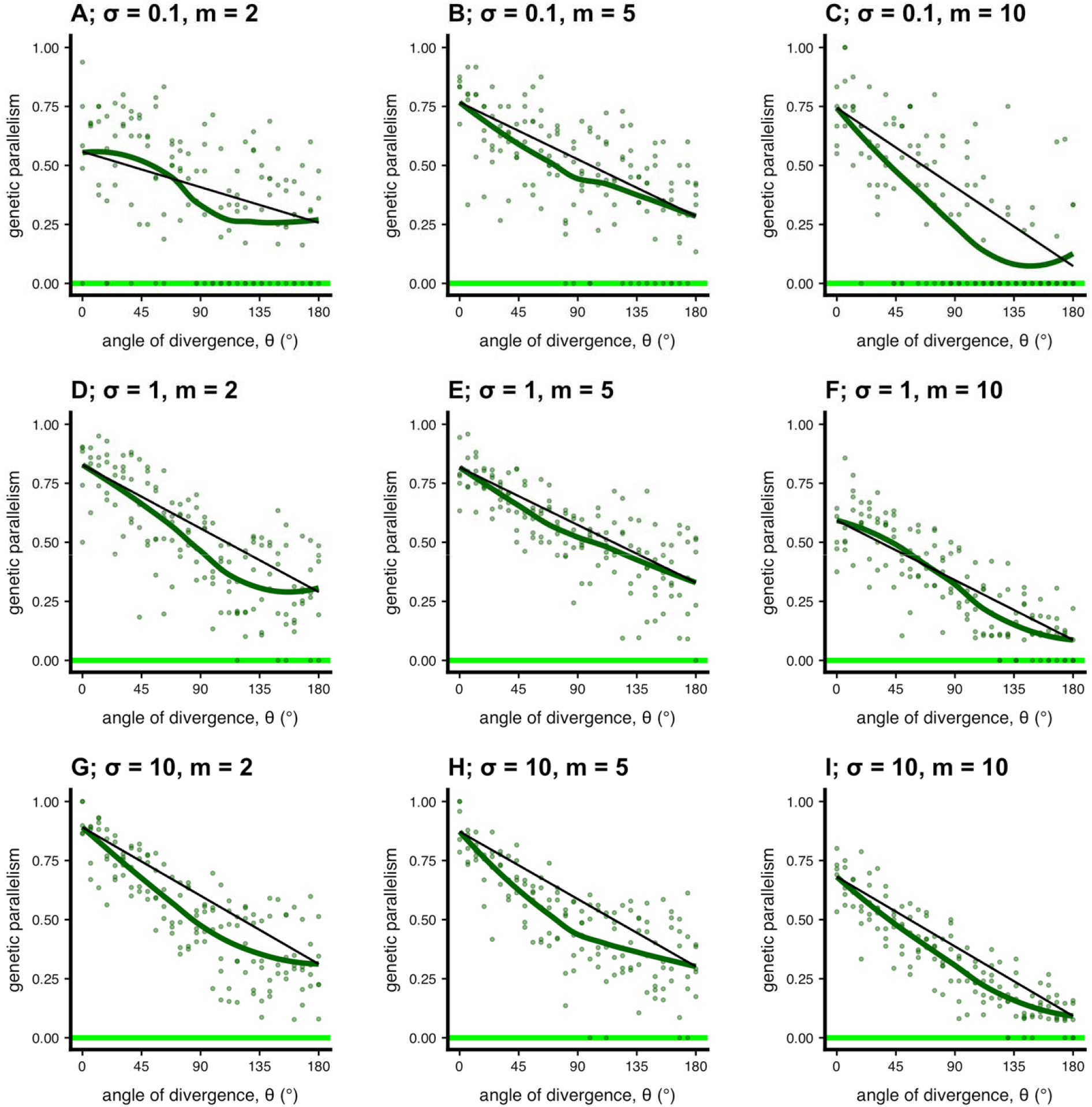
Genetic parallelism across the continuum of parallel to divergent natural selection (pop size = 5000). This figure presents simulations similar to Fig. 2A in the main text but with varying parameter values (selection [*σ*] and dimensionality [*m*]). We ran these particular simulations for *T* = 1000 generations. All other parameters as in main text.

**Figure S4.**
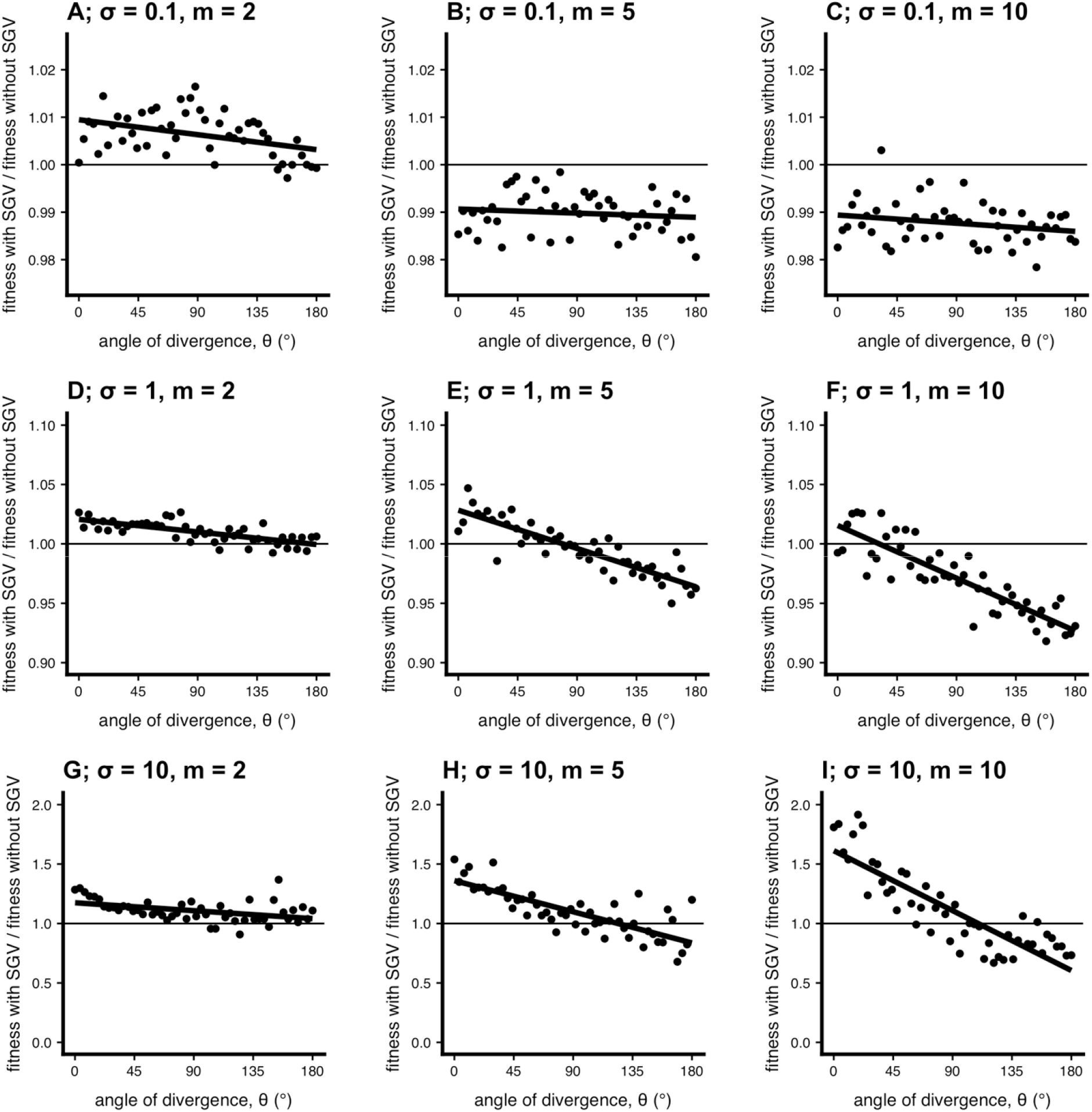
Effect of standing genetic variation on hybrid fitness the continuum of parallel to divergent natural selection (pop size = 100). This figure presents simulations similar to Fig. 3B in the main text but with varying parameter values (selection [*σ*] and dimensionality [*m*]). We ran these particular simulations for *T* = 5000 generations. All other parameters as in main text. Note different y-axis scales across rows.

**Figure S5.**
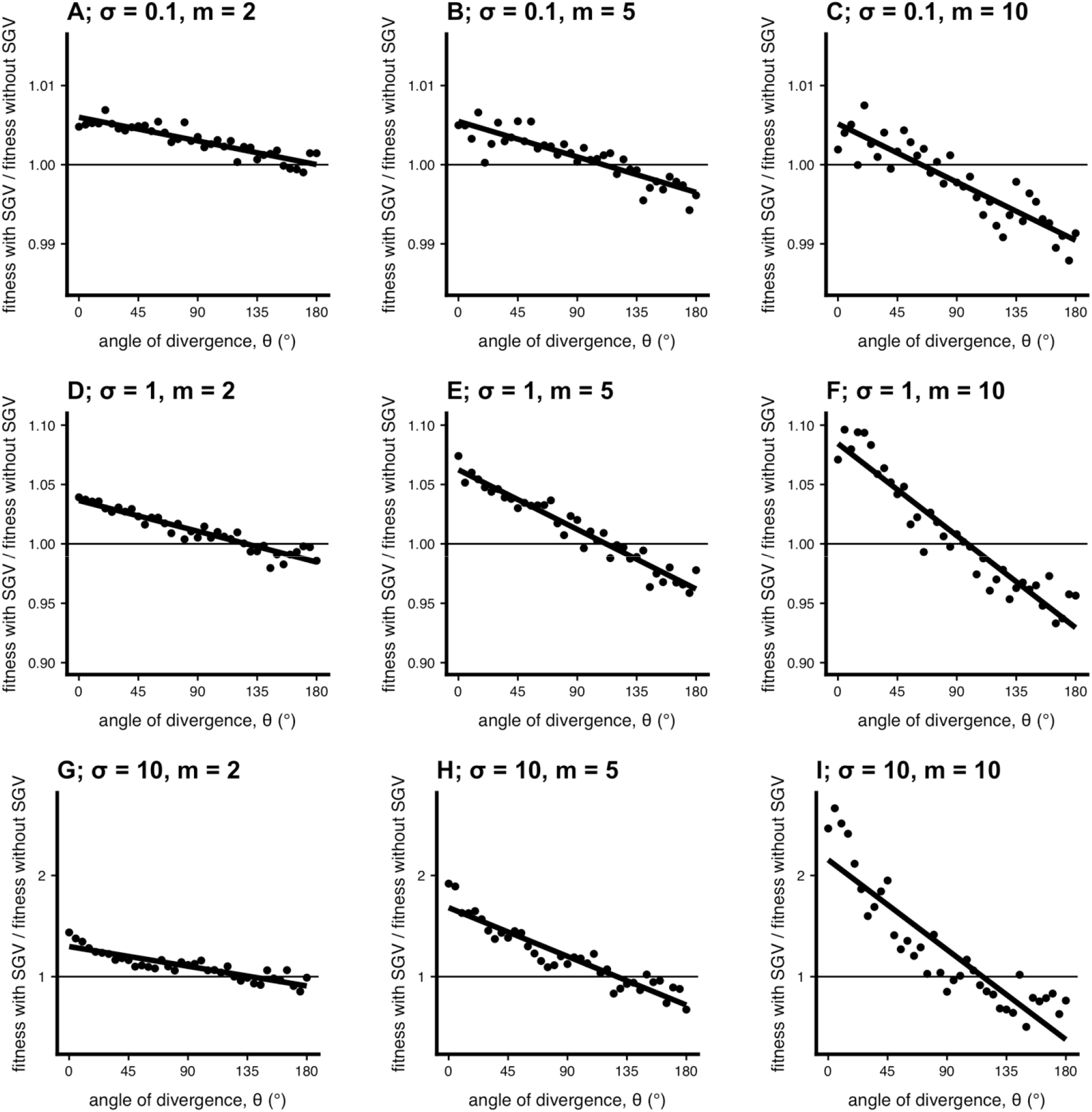
Effect of standing genetic variation on hybrid fitness the continuum of parallel to divergent natural selection (pop size = 1000). This figure presents simulations similar to Fig. 3B in the main text but with varying parameter values (selection [*˃*] and dimensionality [*m*]). We ran these particular simulations for *T* = 2000 generations. All other parameters as in main text. Note different y-axis scales across rows.

**Figure S6.**
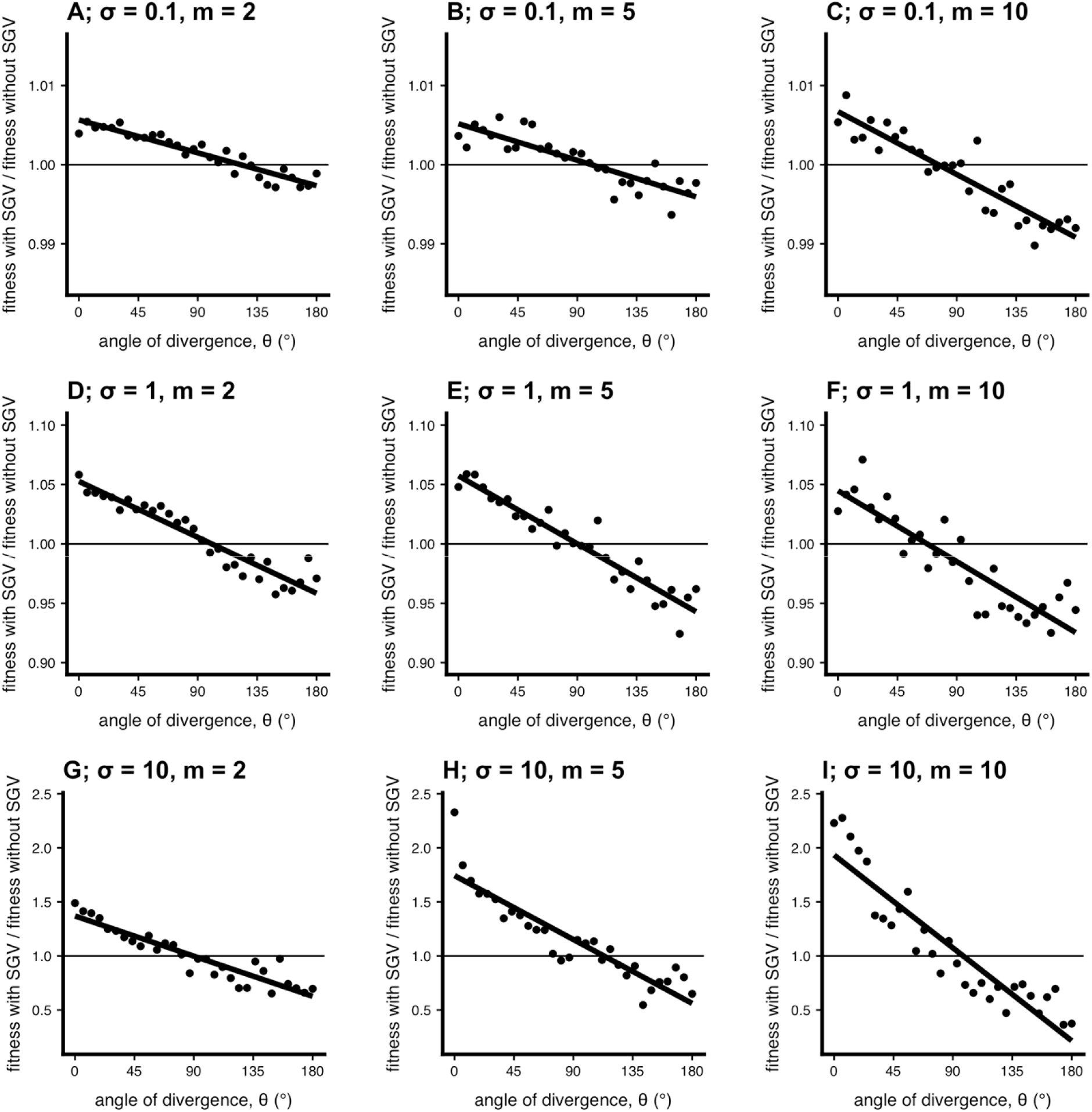
Effect of standing genetic variation on hybrid fitness the continuum of parallel to divergent natural selection (pop size = 5000). This figure presents simulations similar to Fig. 3B in the main text but with varying parameter values (selection [*σ*] and dimensionality [*m*]). We ran these particular simulations for *T* = 1000 generations. All other parameters as in main text. Note different y-axis scales across rows.

**Figure S7.**
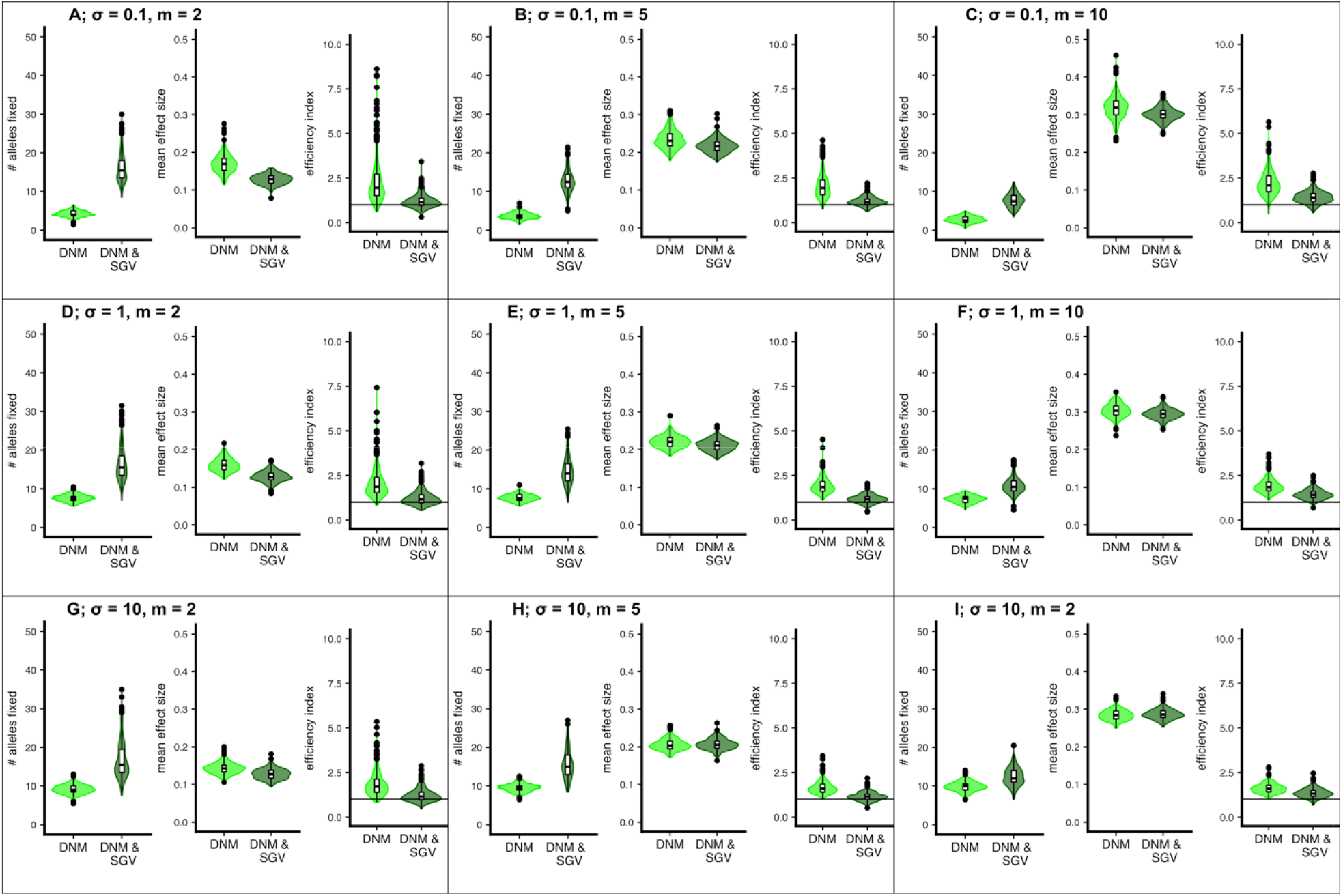
Properties of fixed mutations under a variety of parameter combinations (*N* = 1000). This figure presents simulations similar to Fig. 5 in the main text but with varying parameter values (selection [*σ*] and dimensionality [*m*]). See main text and panel description of Fig. 5 for more detail. Patterns were similar for other population sizes.

**Figure S8.**
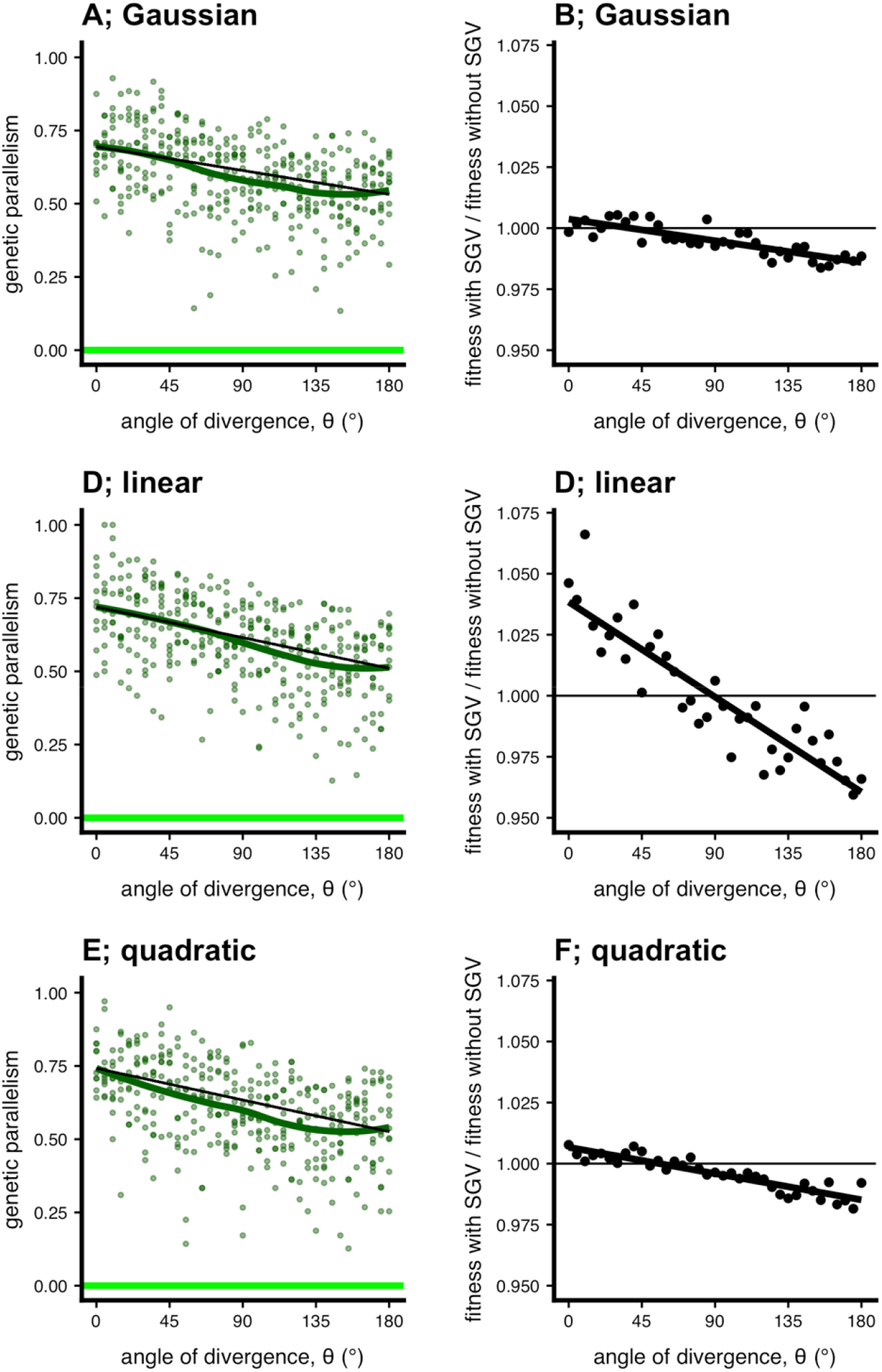
Simulations under various fitness functions. Here we plot simulations across environments for (A & B) Gaussian (*W* = exp(−*σ* ||***z*** − **o**||^2^/2); equation 1), (C & D) linear (*W* = 1 − *σ* ||***z*** − **o**||), and (E & F) quadratic (*W* = 1 − *σ* ||***z*** − **o**||^2^/2) fitness functions. We show results for both genetic parallelism and the effect of standing variation on hybrid fitness. We ran these simulations with a nearer optimum and weaker selection (*d* = 0.5, *σ* = 0.5, *N* = 1000, *m* = 5) because populations otherwise became extinct with linear/quadratic fitness functions. Under these conditions, the non-linear decrease in parallelism is less substantial for all parameter values. Nevertheless, the patterns are qualitatively similar among the three sets of simulations (note differences in *y*-axis scales).

**Figure S9.**
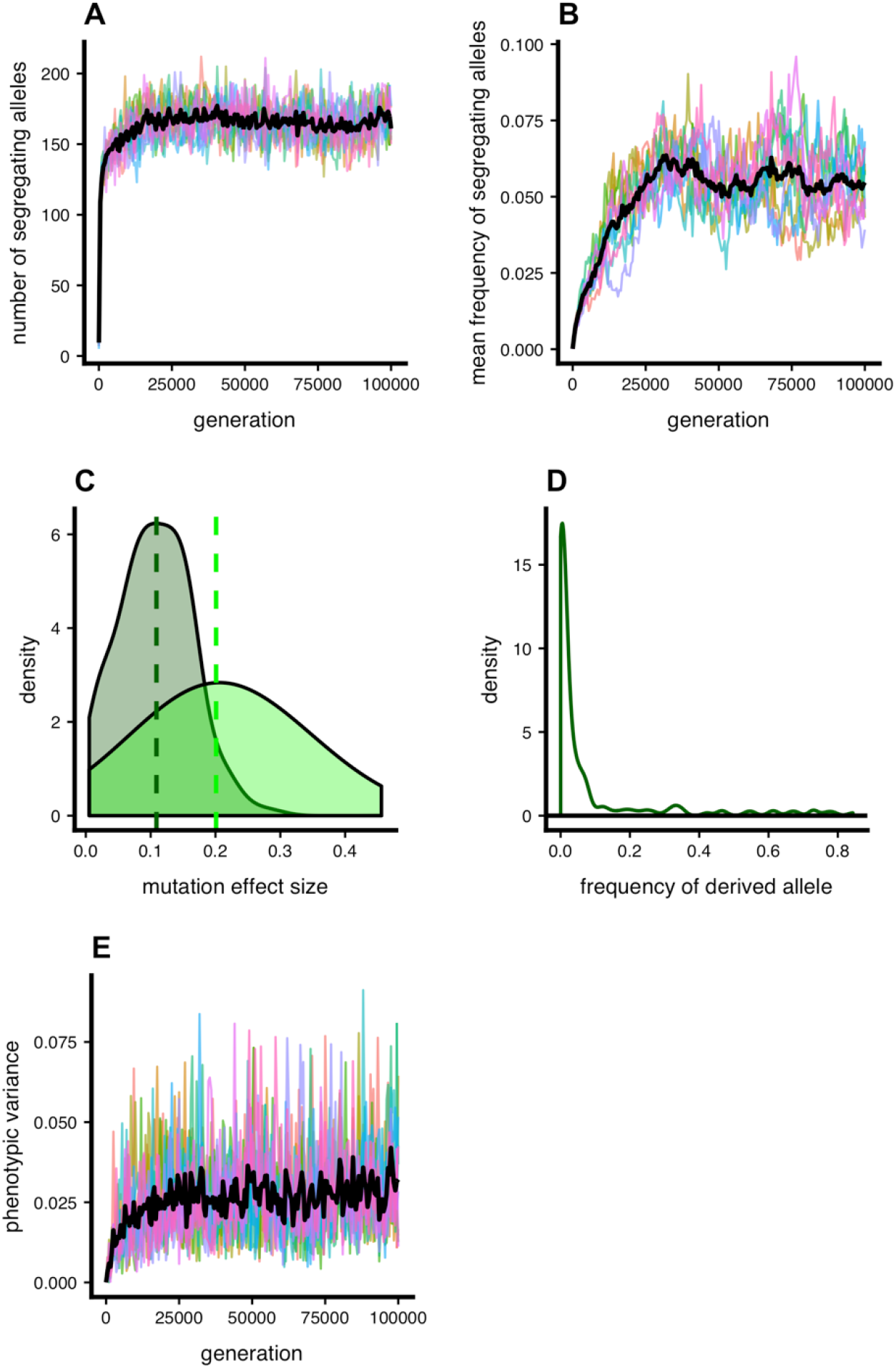
Mutation-selection balance and mutation effect sizes in ancestral populations. In panel (a) we are showing the number of segregating sites in each of 10 ancestral populations and (b) the mean frequency of the derived alleles at each of these sites in the ancestral populations. The black line is plotted through the mean of all populations at each generation, and all ten burnins used to generate our main text results are shown. Panel (c) illustrates the distribution of mutation effect sizes—the Euclidean distance of a mutational vector in phenotypic space—at the end of a single representative burn-in simulation (dark green), as compared to the distribution of mutations that arise *de novo* (light green). The vertical lines represent the median mutation effect size for each group. Panel (d) represents the site-frequency spectrum for segregating sites (excluding sites that have fixed). And panel (e) shows the phenotypic variance in the ancestral population over time. (*m* = 5 for all simulations shown; for rest of parameters see Table 1).

**Figure S10.**
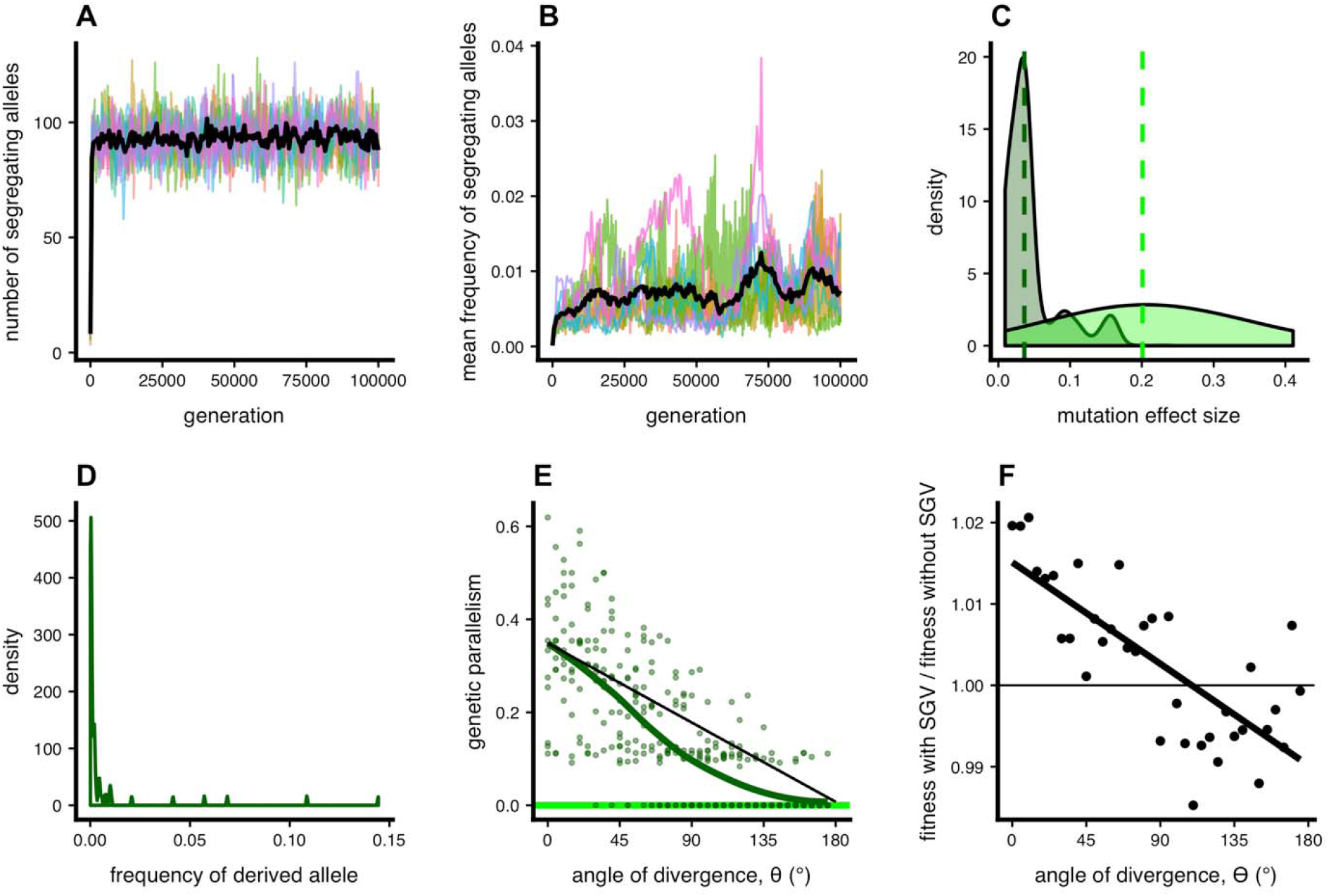
Mutation-selection balance and mutation effect sizes in ancestral populations under stronger selection (*σ*_anc_ = 1). These parameter values imply *μ* << *α*^2^ *α*, as in the House-of-Cards regime (Turelli 1984, 1985) from a Gaussian regime under the alternative set of paremters.. (a) The number of segregating sites in each of 10 ancestral populations and (b) the mean frequency of derived alleles at each of these sites in the ancestral populations. The black line is plotted through the mean of all populations at each generation, and all ten burn-ins used to generate the results ([e] and [f]) are shown. Panel (c) illustrates the distribution of mutation effect sizes—the absolute value of a mutation’s effect on the phenotype—at the end of a single burn-in simulation, as compared to the distribution of mutations that arise *de novo.* The vertical lines represent the median mutation effect size for each group. Panel (d) represents the site-frequency spectrum histogram for segregating sites. (Compare these to Fig. S1). Panels (e) and (f) are as in Fig. 2A and 3B in the main text. For unspecified parameters see Table 1 in the main text. This parameter combination t

**Figure S11.**
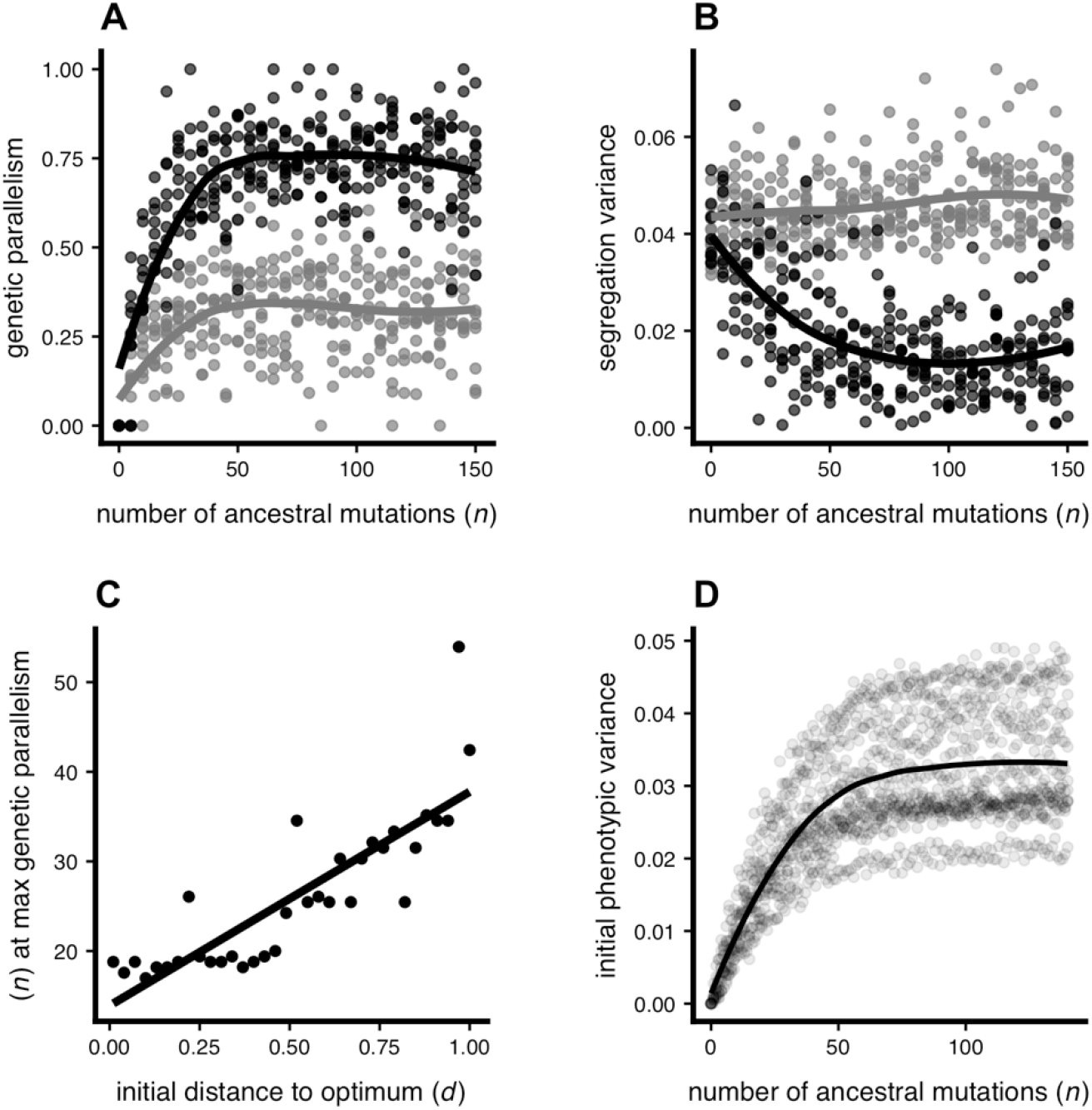
The effects of standing genetic variation on genetic parallelism and phenotypic segregation variance in hybrids under parallel and divergent natural selection. We show (a) genetic parallelism (main text equation 2) and (b) net segregation variance for populations founded with varying quantities of ancestral standing variation (*n*: number of ancestral mutations). Populations were subject to either parallel (*θ* = 0°; black) and divergent (*θ* = 180°; grey) selection, with d=1, and there were 10 replicate simulations per parameter combination. Genetic parallelism values of 0 indicate no parallelism and values of 1 indicate complete parallelism (main text Eq. 2). The curves are loess fits. Panel (c) shows that the quantity of ancestral standing variation that maximizes genetic parallelism under parallel selection (= 0°) increases when populations adapt to more distant optima. A value of *d* = 1 is 10 mutational SDs. The line is a linear regression. Panel (d) shows the relationship between the genetic (phenotypic) variation in a parental population as a function of *n*.

**Figure S12.**
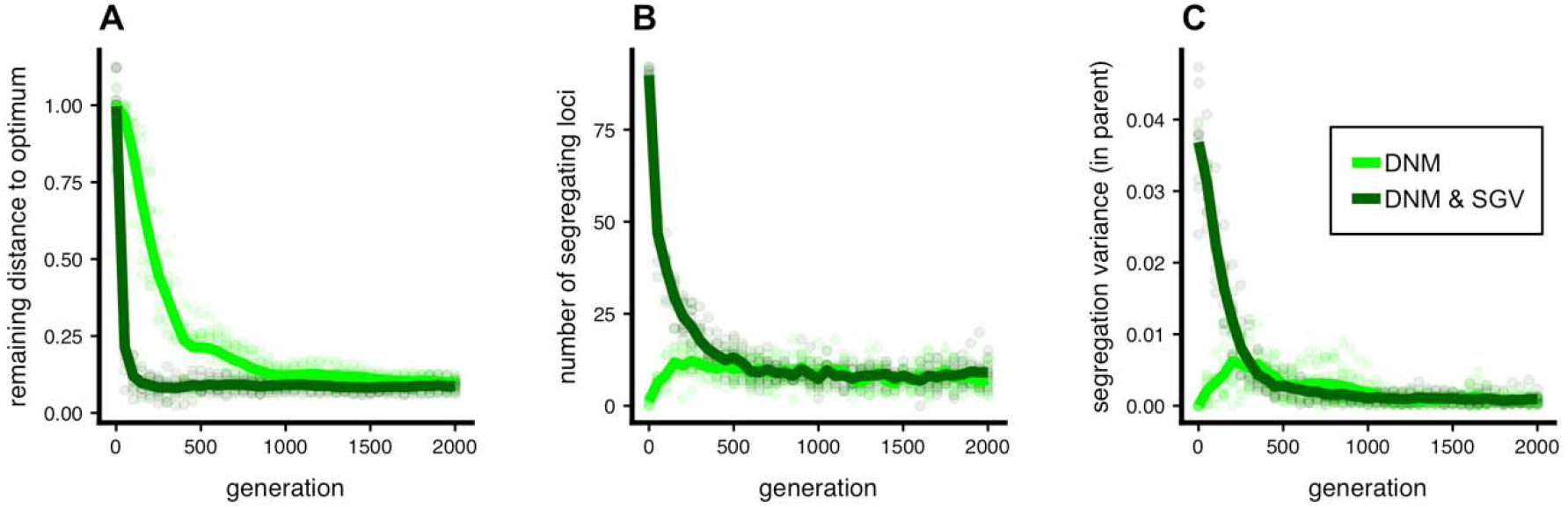
Effect of standing variation on the pace of adaptation and attainment of mutation-selection-drift balance. (a) Populations that adapt with standing variation in addition to new mutation (DNM & SGV; *n* = 100 segregating alleles; dark green) reach the phenotypic optimum more quickly than populations that adapt from new mutation only (DNM; *n* = 0 segregating alleles; light green). (b) Although populations equipped with standing variation adapt more quickly than populations adapting from new mutation only, they both reach mutation-selection-drift balance by generation 2000. (c) The phenotypic (genotypic) variance in parental populations, calculated as it is in hybrids (see main text), is stable and near zero by the end of each simulation. The initial distance to the optima, *d,* is 1 for all simulations. We plot 10 replicate simulations, and lines connect the mean values at each sampled generation. For unspecified parameters see Table 1 in the main text.

**Figure S13.**
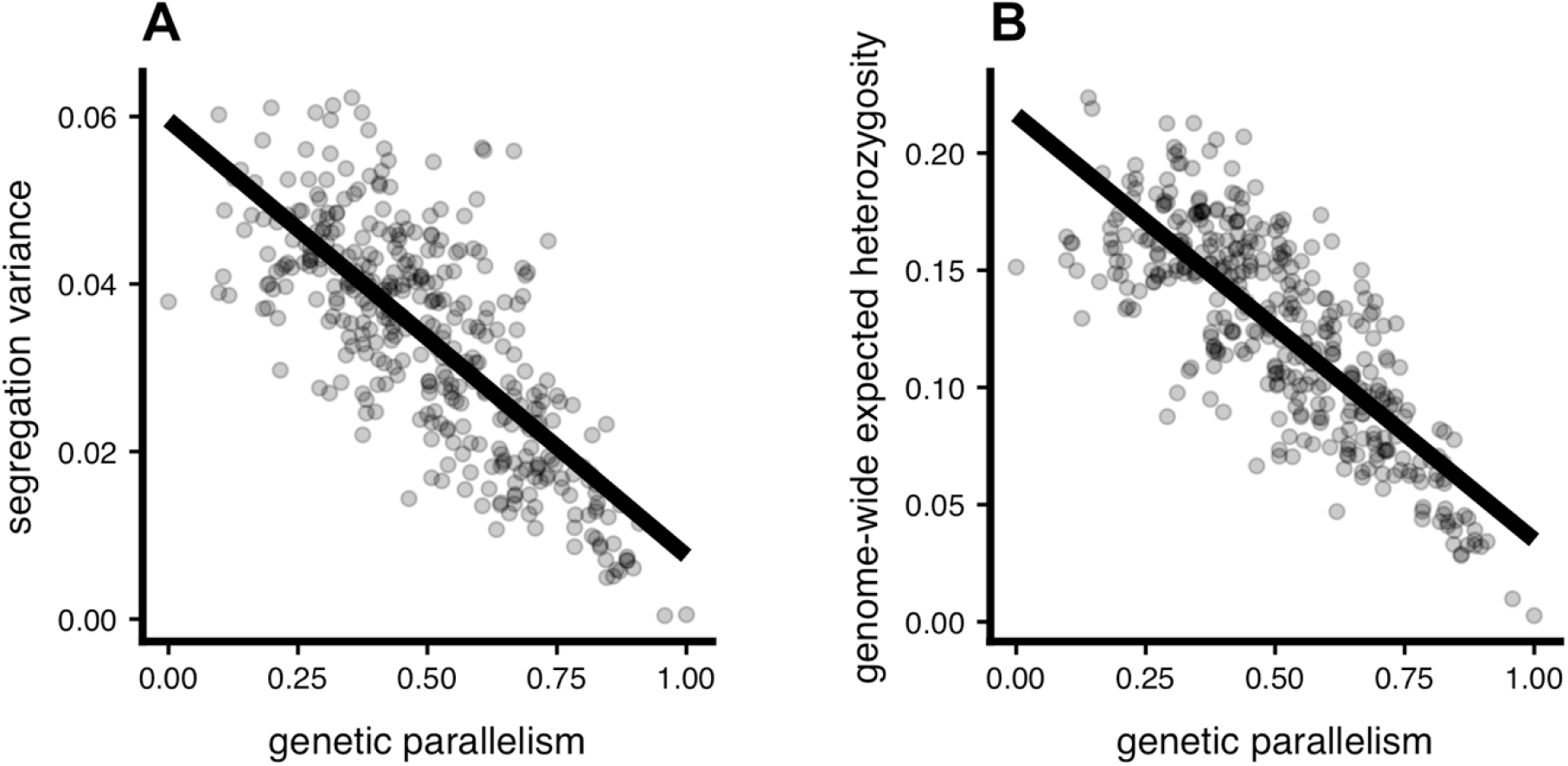
Relationship between genetic parallelism and (a) segregation variance and (b) expected heterozygosity. Our metric of genetic parallelism (main text equation 2) is on the *x*-axis. This is the data plotted in Fig. 2A & 2C of the main text. We show the correlation between genetic parallelism and (a) segregation variance (*r*^2^ = 0.56) and (b) genome-wide expected heterozygosity (2*p*[1-*p*], averaged across all loci (*r*^2^ = 0.63). Patterns were similar for F_ST_ (Hudson et al. 1992) and net *π* (Nei and Li 1979) (not shown).

**Figure S14.**
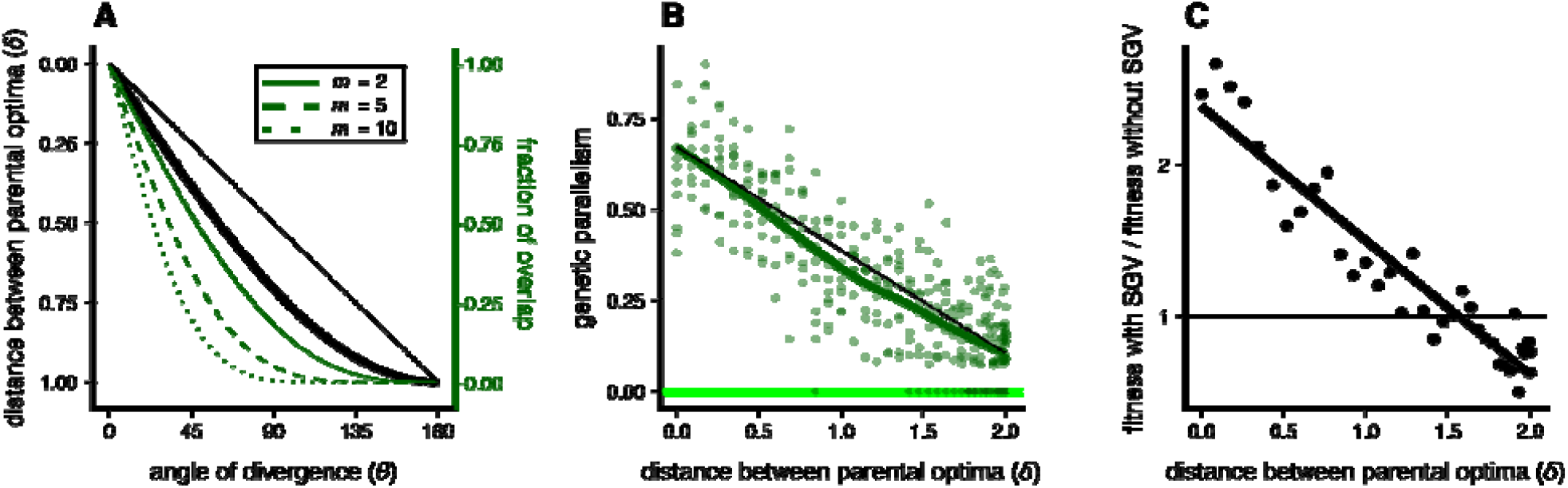
Alternative presentation of simulation results across environments: distance between optima ( ). Panel (a) plots the relationship between the angle of divergence, *θ,* and the Euclidean distance between parental optima, (thick black line; note reversal of y-axis; scaled between 0 and 1 by dividing by 2*d*). We also plot the fraction of non-overlap as in the main text Fig. 2B (thin dark green lines; *m* = 2, 5, 10 in increasing steepness). Panel (b) shows genetic parallelism vs.. For a given value of is invariant with dimensionality (i.e., the distance between optima does not change as dimensionality increases). Accordingly, the nonlinearity emerges even when considering, but only appreciably when considering higher dimensions (*m* > 5). In both panels, the thin and straight black line connects the fit at 0° with 180° for visual reference. In panel (c) we show the results for hybrid fitness, similar to Fig. 3B in the main text. All panels show simulations conducted in 10 dimensions (*m* = 10, *=* 1, *N* = 1000).

**Figure S15.**
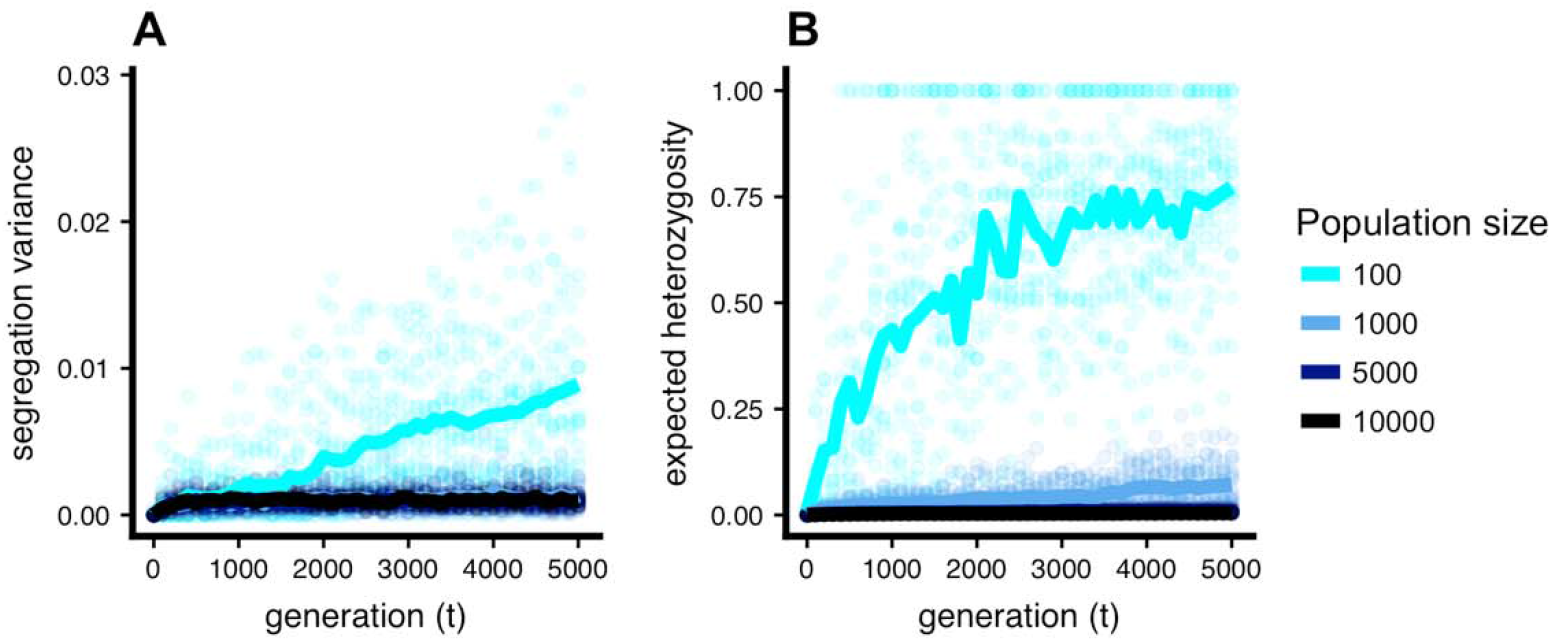
The effect of population size on the rate of divergence between populations due to drift. We show populations held at a common optimum with no standing variation (i.e. *d* = 0, *n* = 0) and plot (a) segregation variance and (b) expected heterozygosity in hybrids over time for 5,000 generations. The evolution of segregation variance is proportional to the rate of evolution of reproductive isolation under parallel natural selection. Greater drift in smaller populations leads to greater segregation variance and heterozygosity. The lines are drawn as the average of 10 replicate simulations (*m* = 5, *σ* = 1).

**Figure S16.**
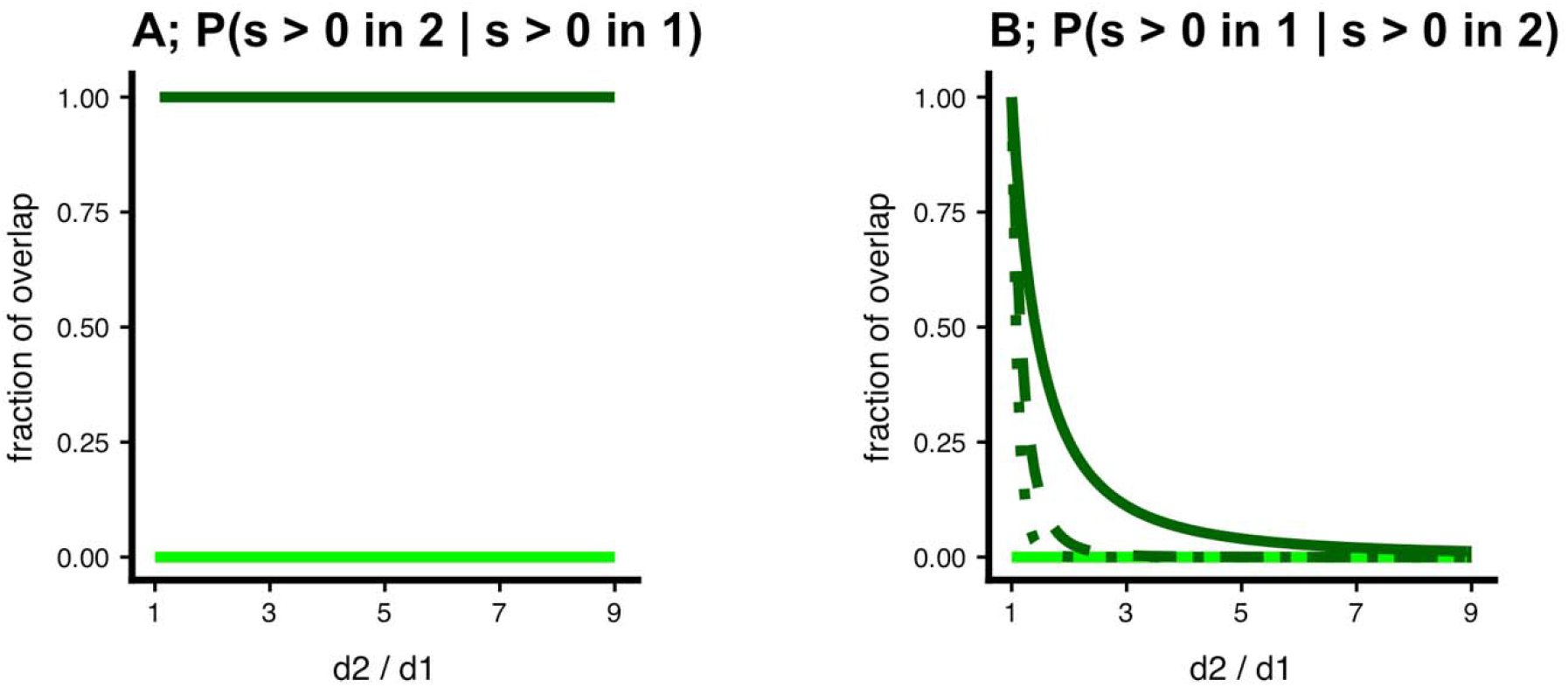
Fraction of overlap of beneficial mutations with parallel selection (*θ* = 0°) but unequal distance (*d*_1_ ≠ *d*_2_). The main text explores how the fraction of overlap changes with theta while holding *d*_1_ = *d*_2_ = *d* constant. Here we explore how the fraction of overlap changes with the ratio d_2/d_1 when holding *θ* = 0° constant. Unlike the metric presented in the main text this metric is asymmetrical because one population is completely contained within the other. Panel (A) plots the fraction of overlap for population 1 (the fraction of alleles that are beneficial in population 1 are also beneficial in population 2) as a function of *d*_2_ / *d*_1_. With *d*_1_ < *d*_2_ the value is 1 for any ratio *d*_2_ / *d*_1_ because population 1’s hypersphere is contained within population 2’s. Panel (B) plots the fraction of alleles that are beneficial in population 2 that are also beneficial in population 1. This latter result mirrors what is seen in the main text Fig. 3C: as the locations of the optima depart from one another the fraction of overlap rapidly approaches zero and does so most rapidly at the onset of departure.

**Figure S17.**
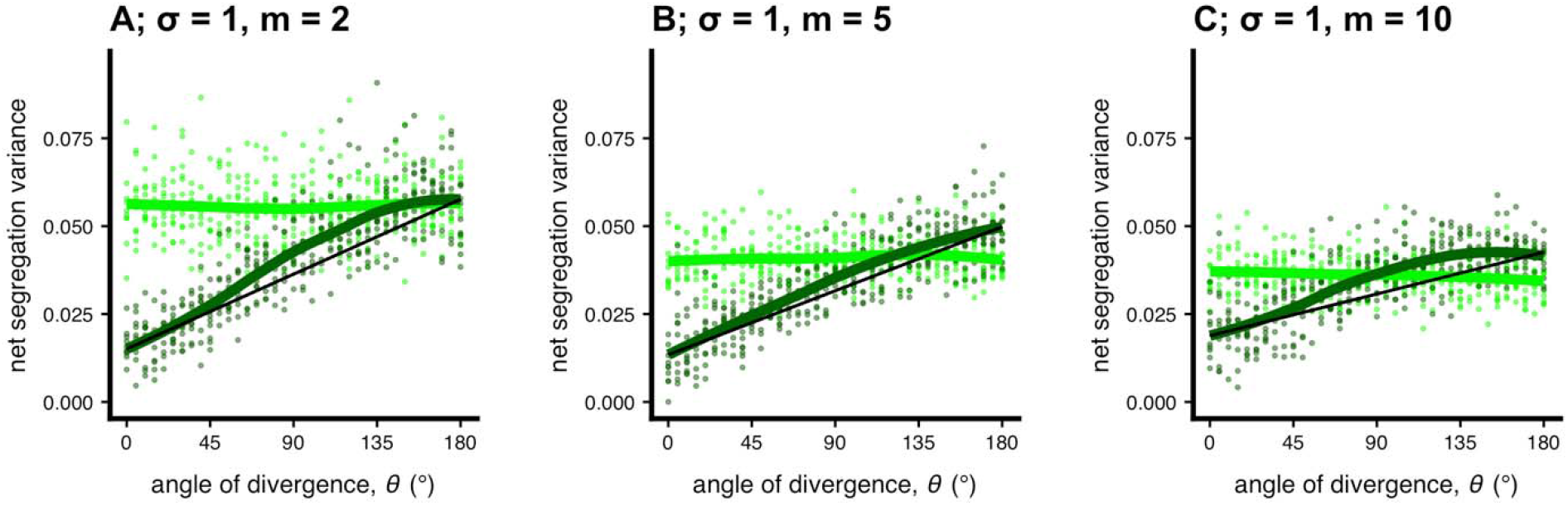
Effect of dimensionality on net segregation variance. These plots are similar to Fig. 2C in the main text except we show results for three different dimensionalities. Under divergent natural selection, standing genetic variance increases segregation variance relative to de novo mutations alone more at higher dimensionalities.

**Figure S18.**
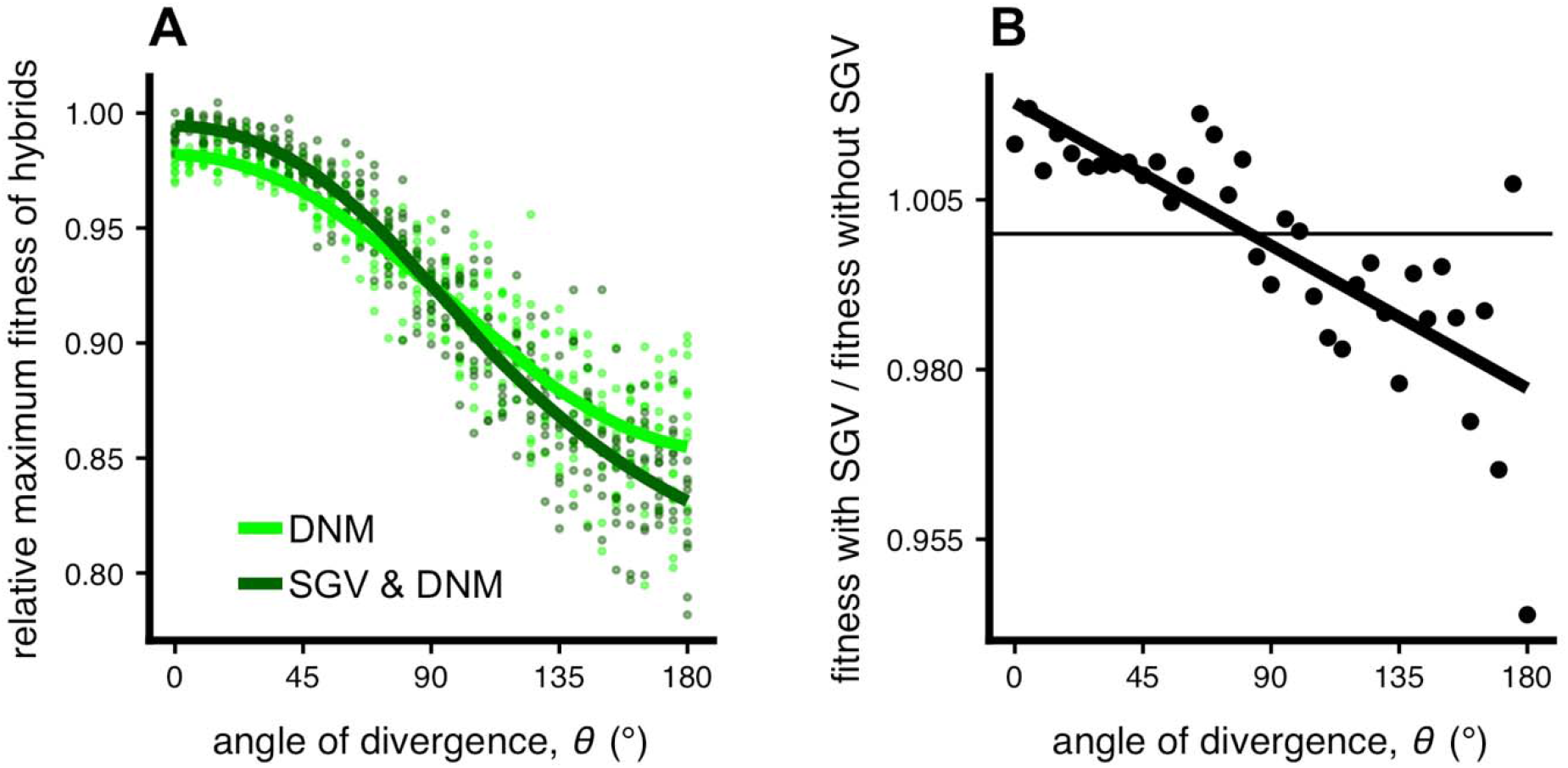
The effect of standing genetic variation (SGV) on relative maximum hybrid fitness across environments. Data are from simulations plotted in the main text, but instead of mean fitness of all hybrids we depict the mean fitness of the top 5 % of hybrids relative to the mean fitness of parents. We plot both the (a) raw values of relative maximum fitness and (b) the effect of standing variation on maximum hybrid fitness (dark green divided by light green).

